# Spliced peptides and cytokine driven changes in the immunopeptidome of melanoma

**DOI:** 10.1101/623223

**Authors:** Pouya Faridi, Katherine Woods, Simone Ostrouska, Cyril Deceneux, Ritchlynn Aranha, Divya Duscharla, Stephen Q. Wong, Weisan Chen, Sri Ramarathinam, Terry C.C. Lim Kam Sian, Nathan P. Croft, Chen Li, Rochelle Ayala, Jonathan Cebon, Anthony W. Purcell, Ralf B. Schittenhelm, Andreas Behren

**Affiliations:** Department of Biochemistry and Molecular Biology and Infection and Immunity Program, Monash Biomedicine Discovery Institute, Monash University, Clayton, Victoria, Australia; Cancer Immunobiology, Olivia Newton-John Cancer Research Institute, Austin Hospital, Heidelberg, Victoria, Australia; School of Cancer Medicine, La Trobe University, Bundoora, Victoria, Australia; Cancer Research Division, Peter MacCallum Cancer Centre, Melbourne, Victoria, Australia; Department of Pathology, Peter MacCallum Cancer Centre, Melbourne, Victoria, Australia; Department of Biochemistry and Genetics, La Trobe Institute for Molecular Science, La Trobe University, Melbourne, Victoria, Australia; Department of Biology, Institute of Molecular Systems Biology, ETH Zürich, Switzerland; Monash Proteomics & Metabolomics Facility, Monash Biomedicine Discovery Institute, Monash University, Clayton, Victoria, Australia

**Author notes:** These authors contributed equally (Pouya Faridi, Katherine Woods).

**Keywords:** Immunopeptidome, Immunoproteasome, Transpeptidation, CD8^+^ T cells, Post-translational splicing, Melanoma, Antigen processing, IFN□, Inflammation, HLA, Immunotherapy

## Abstract

Antigen-recognition by CD8^+^ T cells is governed by the pool of peptide antigens presented on the cell surface in the context of HLA class I complexes. Recent studies have shown not only a high degree of plasticity in the immunopeptidome, but also that a considerable fraction of all presented peptides is generated through proteasome-mediated splicing of non-contiguous regions of proteins to form novel peptide antigens. Here we used high-resolution mass-spectrometry combined with new bioinformatic approaches to characterize the immunopeptidome of melanoma cells in the presence or absence of interferon-γ. In total, we identified more than 60,000 peptides from a single patient derived cell line (LM-MEL-44) and demonstrated that interferon-γ induced marked changes in the peptidome with an overlap of only ∼50% between basal and treated cells. Around 6-8% of the peptides were identified as *cis*-spliced peptides, and 2213 peptides (1827 linear, 386 *cis*-spliced peptides) were derived from known melanoma-associated antigens. These peptide antigens were equally distributed between the constitutive and interferon-γ induced peptidome. We next examined additional HLA-matched patient derived cell lines to investigate how frequently these peptides were identified and found that a high proportion of both linear and spliced peptides were conserved between individual patient tumors, drawing on data amassing to over 100,000 peptide sequences from these extended data sets. Moreover, several of these peptides showed *in vitro* immunogenicity across multiple melanoma patients. These observations highlight the breadth and complexity of the repertoire of immunogenic peptides that can be exploited therapeutically and suggest that spliced peptides are a major new class of tumor antigens.

## Introduction

Antigen recognition by cytotoxic T cells and subsequent tumor cell destruction is the key component underlying cancer immunotherapy strategies. Its importance has been widely demonstrated, and loss-of-function of elements in the antigen processing and presentation pathways has been shown to confer therapeutic resistance(1). Correlative findings point to neo-antigens arising from tumor mutations as an important source of immunogenic antigens in the context of melanoma and other cancers(2). Nonetheless, tumors with low mutational burdens can respond to checkpoint inhibitor therapy, and the presence of a high tumor-mutational load does not necessarily correspond to the efficacy of treatment(3). The human leukocyte antigen (HLA) class I-bound peptides (*p*-HLA-I) arising from the mutant proteins are mostly heterogeneously expressed and additionally determined by the patient-specific HLA-subtypes, making predictions about their presentation and immunogenicity unreliable. While a number of recent studies have reported the utility of mass spectrometry combined with exome sequencing in identifying HLA-presented peptides derived from mutated proteins(4–6), the analysis of the contribution of mutational neoantigens to the overall tumor immunogenicity remains complicated and unresolved.

Against this background, the composition of the immunopeptidome, or the repertoire of HLA-bound peptides presented on the surface of the cell and their contribution to tumor immune recognition, becomes significant. The immunopeptidome is largely shaped by antigen processing through the proteasome complex for subsequent presentation of short peptide epitopes on MHC molecules(7). Several forms of the proteasome complex exist, each with differing enzymatic activities(8). In melanoma cells, the constitutive proteasome is expressed under steady state conditions. The expression of an immunoproteasome, the subtype expressed by dendritic cells and other cells of the immune system, may be induced in tumor cells in a cytokine-dependant manner (9), leading ultimately to changes in the peptides presented to the immune system(10, 11). We have previously demonstrated induction of the immunoproteasome in a range of human melanoma cell lines in the presence of the inflammatory cytokine IFN□ *in vitro*, and in melanoma patient inflamed tumors (characterized by presence of tumor infiltrating lymphocytes (TILs)) *ex vivo*(12). Dependant on the proteasome subtype expressed by the cell, we have shown that a single melanoma antigen (NY-ESO-1) can be processed into several different epitopes. These differences in antigen processing led to concomitant change in the ability of antigen specific T cells to target the tumor cell. Thus, the potential for a tumor cell to ‘look’ substantially different to CD8^+^ T cells, depending on inflammation at the tumor site, arises. Moreover, recent studies by ourselves and others(13, 14) have shown that a significant proportion of *p*-HLA-I are not genomically templated and result from post-translational proteasome splicing (ligation of non-contiguous small polypeptide segments from the same or different proteins). To date, these peptides have been missed in most neoantigen discovery studies due to the lack of appropriate bioinformatics tools(15, 16).

In this study we have used high resolution mass spectrometry approaches combined with a novel bioinformatics workflow to identify linear and spliced *p*-HLA-I presented in the melanoma immunopeptidome in the presence or absence of the cytokine IFNγ. These included a number of linear and *cis*-spliced peptides derived from melanoma–associated antigens. A series of identified linear and *cis*-spliced peptides were tested for *in vitro* immunogenicity across multiple melanoma patients and healthy donors. While the peptide repertoire changed significantly, immunogenicity of selected peptide pools from different treatment conditions (+/- IFNγ) *in vitro* did not change across melanoma patients. However, T lymphocyte responses to pools of IFNγ upregulated peptides were not seen in healthy donors. We also demonstrate that *cis*-spliced peptides were widely presented by melanoma cells and immunogenic in multiple donors. These findings have significant implications for cancer immunotherapy as well as for fundamental questions such as induction of immune-tolerance, T cell repertoires and immune recognition.

## Results

### The melanoma immunopeptidome is composed of linear and spliced peptides

We have established a comprehensive repository of HLA class I peptide ligands presented by a patient derived melanoma cell line (LM-MEL-44) utilizing either the constitutive proteasome (-IFN-γ) or the immunoproteasome (+IFN-γ). Using three biological replicates for each condition, we identified around 60,000 peptides presented across all HLA class I allotypes expressed by LM-MEL-44 cells (Table S1). Approximately 6-8% of the peptides in each sample were conservatively assigned as *cis*-spliced in origin (Fig. 1A, Table S1), being derived from non-contiguous sequences of the same protein. This proportion of peptides of *cis*-spliced origin is in agreement with previous studies (14,17,18). As expected for HLA class I epitopes, the majority of peptides were 9 amino acids in length with no apparent difference between linear and *cis*-spliced sequences (Fig.1B).

**Figure 1.**
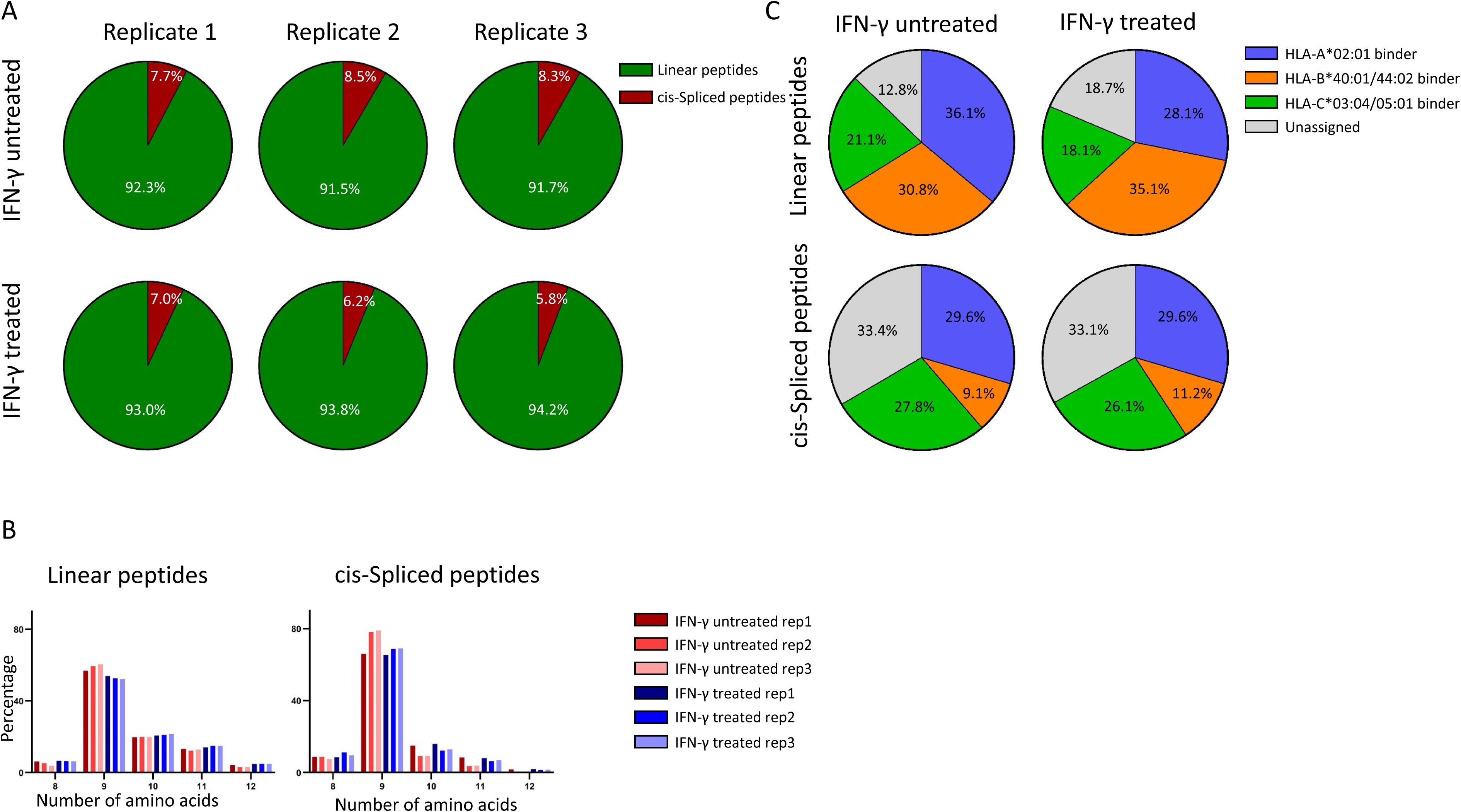
**The melanoma immunopeptidome consists of linear and spliced peptides in the presence and absence of IFN**D**. A**, 6-8 % of peptides presented by the LM-MEL-44 cell line were *cis*-spliced peptides across 3 replicates. **B**, Length distribution analysis showed that majority of linear and *cis*-spliced peptides were 9 mers in the presence and absence of IFN□ **C**, Using the NetMHC4.0 algorithm, binding of identified 8-12 mer peptides to all HLA-I haplotypes presented by LM-MEL-44 was predicted.

Using NetMHC 4.0 binding prediction algorithm, more than 80% of the linear peptides were assigned to at least one of the HLA class I alleles expressed on the surface of the LM-MEL-44 cells (HLA-A*02:01, B*40:01/*44:02, C*03:04/*05:01)(19) suggesting that the majority of the identified peptide sequences can be considered genuine HLA class I ligands (Fig. 1C, Table S2). The percentage of *cis*-spliced peptides predicted to bind to HLA-A or HLA-C molecules was found to be comparable to linear epitopes, but intriguingly substantially fewer *cis*-spliced peptides were predicted to bind to HLA-B*40:01 and HLA-B*44:02, suggesting that LM-MEL-44 cells generate a lower number of *cis*-spliced peptides that conform to the consensus-binding motif of these HLA-B allotypes (Fig. 1C). Moreover, a significantly higher percentage of unassigned sequences was observed amongst the *cis*-spliced epitopes, which is in agreement with previous reports (13, 14) and which can be attributed to the fact that binding algorithms such as NetMHC 4.0 are exclusively trained on linear peptide sequences.

In addition, it should be noted that we did not identify in our entire dataset any of the mutational neoantigens that have been described for the LM-MEL-44 cell line based on exome sequencing data (20). However, this is not surprising as this cell line has a relatively low mutational load (Table S3).

### The generation of spliced peptides is not a random process

To address whether spliced peptides are randomly generated, we comparatively analyzed the overlap of both linear and spliced peptides across our three biological replicates. A total of 1399 and 1795 *cis*-spliced peptides were identified in at least two of the three biological replicates for the IFN□ -untreated and treated samples, which corresponds to 47.6% and 52.2%, respectively (Fig. 2A). Importantly, a similar overlap was observed for the linear epitopes (48.8% and 53.9%, respectively), which suggests that the generation of *cis*-spliced peptides is not a random process. Of note, the comparatively low overlap between the replicates can be rather attributed to the stochastic nature of data-dependent acquisition mass spectrometry (DDA-MS), which is particularly pronounced when acquiring highly complex samples that contain individual analytes of low abundance (such as HLA peptide samples).

**Figure 2.**
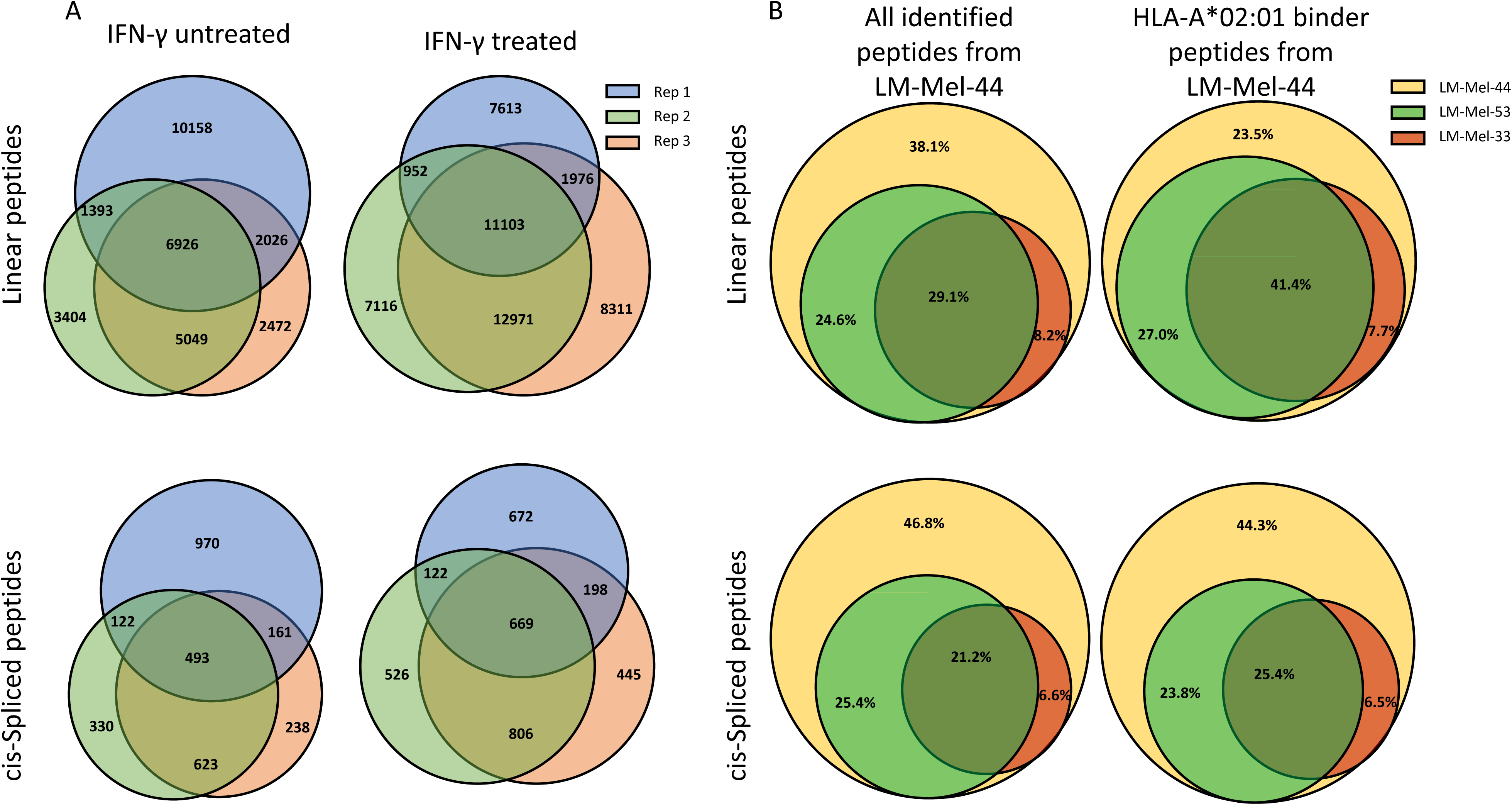
**Generation and presentation of spliced peptides is not random process**. **A,** Reproducibility of linear and *cis*-spliced peptides across biological replicates with around half of the of linear *cis*-spliced of peptides identified in at least two out of three replicates. **B,** A high proportion of linear and spliced peptides presented on LM-MEL-44 cell were also identified on other melanoma cell lines. The yellow circle (defined as 100%) represents all peptides that were identified across all six LM-MEL-44 samples. The green and orange circles represent the proportion of those peptides that were also identified to be presented on LM-Mel-53 and LM-Mel-33, respectively.

To investigate whether the identified *cis*-spliced peptides are also expressed on other cell lines with a similar HLA signature, we analysed the cell lines LM-MEL-53 and LM-MEL-33 by DDA-MS. LM-MEL-53 cells are derived from the same patient as LM-MEL-44 cells, but from another metastasis at a different point in time (21). In contrast, LM-MEL-33 (HLA-A*02:01/A*03:01, B*40:02/*47, C*03:04/*06:02) cells have been isolated from a different patient that shares three HLA alleles with LM-MEL-44 (21). 47% and 28% of the identified *cis*-spliced peptides from LM-MEL-44 were also identified on LM-MEL-53 and LM-MEL-33 cells, respectively (Fig. 2B, Table S1), which further confirms that the generation of *cis*-spliced peptides is not a random process, but more importantly, that *cis*-spliced peptides have a significant potential for cancer immunotherapy.

### IFNγ-treatment alters melanoma HLA class I immunopeptidome

Considering the well-described clinical relevance of so called “hot” versus “cold” tumor microenvironments and previous work demonstrating the influence of cytokine exposure on antigen-presentation pathways (10), we wanted to examine the impact of IFNγ exposure on the immunopeptidome. IFNγ treatment led to the identification of a considerably higher number of HLA epitopes than that from untreated cells, consistent with the upregulation of HLA molecules at the cell surface (Fig. 3A, Fig. S1) (10). Moreover, for common peptides we observed, on average, an increase in peptide abundance in the cytokine treated samples. Of note, only 44.7% of the linear and 52.5% of the spliced peptides were identified under both conditions suggesting that the addition of IFN□ significantly impacts the composition of the immunopeptidome.

**Figure 3.**
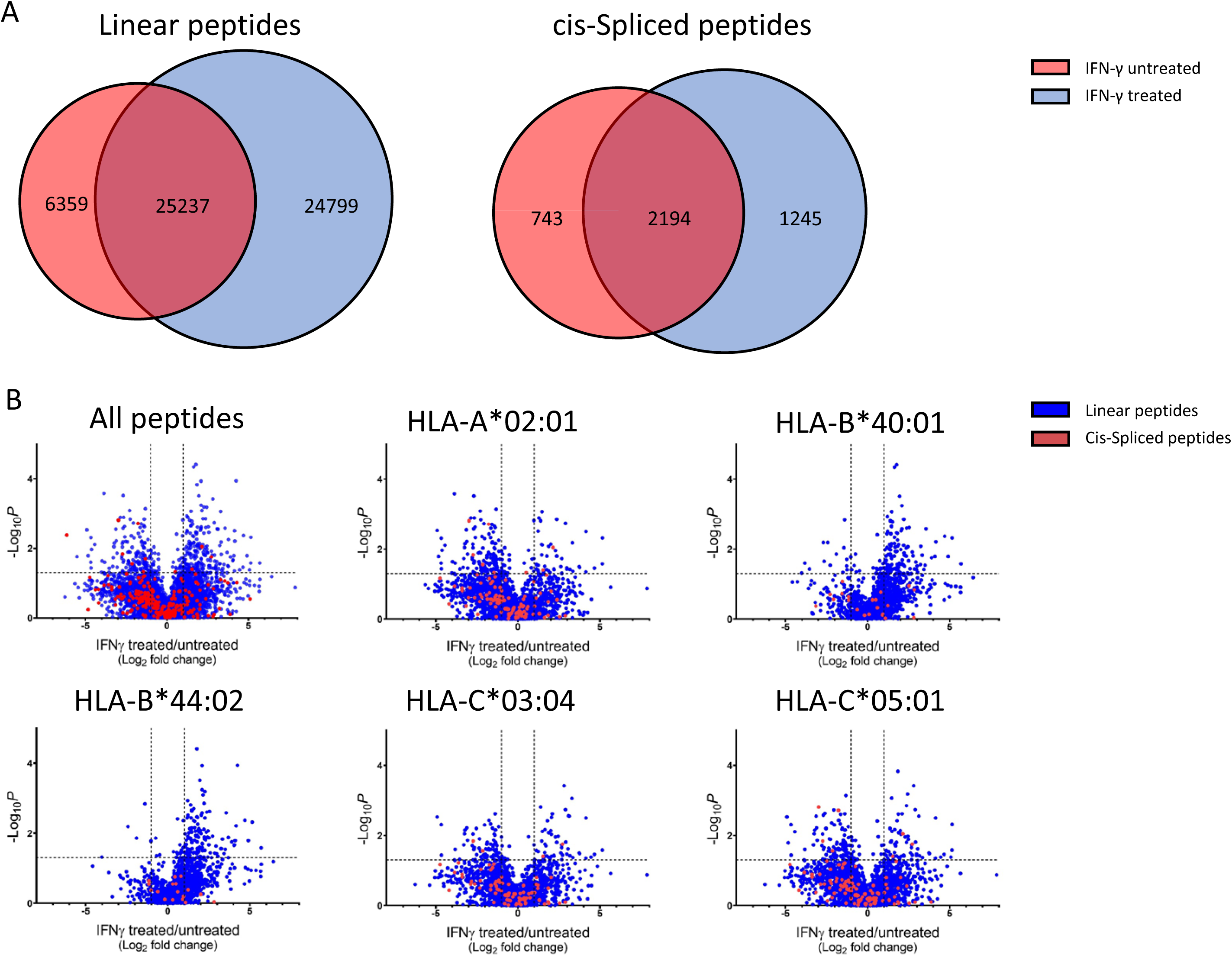
**The melanoma immunopeptidome is substantially altered following exposure to IFN**γ**. A,** Around 20% of linear and 25% of spliced peptides presented by HLA-I were lost following treatment with IFN□, alongside a concurrent increase in novel peptides. **B,** Among peptides identified in 3 replicates both with and without IFN□, an elevation in the number of peptides that bound to HLA-B, compared to HLA-A and C, was observed.

To understand whether IFNγ exposure changes the abundance of individual epitopes independent of the HLA expression levels, we identified epitopes that were present across all replicates (a total of 4942 peptides) and calculated their log2 fold change between IFN□ treated and untreated samples after median normalization of their ms1 intensities to remove any bias introduced through varying levels of IFNγ -induced HLA expression (Fig. 3B, Table S1). A considerable number of epitopes changed in abundance by a factor of at least 2 (both up- and down), confirming that the addition of IFN□ substantially alters HLA class I presentation, while not affecting the proportion of presented *cis*-spliced epitopes on LM-MEL-44 melanoma cells. Interestingly and despite median-normalized ms1 intensities, most of the peptides predicted to bind to HLA-B*40:01 and HLA-B*44:02 were still upregulated upon IFN□ exposure, which correlates to the enhanced upregulation of HLA-B molecules in response to IFN□ compared to HLA-A and HLA-C molecules (10).

### Identification of novel cancer specific peptides in the melanoma immunopeptidome

Next we screened a panel of identified peptides including sequences from melanoma/cancer-associated antigens (MAA) (22, 23) and tumor antigens with demonstrated immunogenicity(24–26). We identified a total of 2213 peptides in our dataset (1827 linear, 386 spliced peptides) derived from 142 different MAAs (Fig. 4A, Table S4). A large proportion (∼45%) of linear peptides have not been previously reported (Fig. 4B) (24). Furthermore, almost all of those peptides generated by splicing constitute potentially novel epitopes. Of the previously reported epitopes, the majority were detected in both the presence and absence of IFN□, whereas >58% of novel peptides were exclusive to IFN□ treated samples, demonstrating the importance of carefully considering experimental conditions for epitope discovery (Fig. 4C).

**Figure 4.**
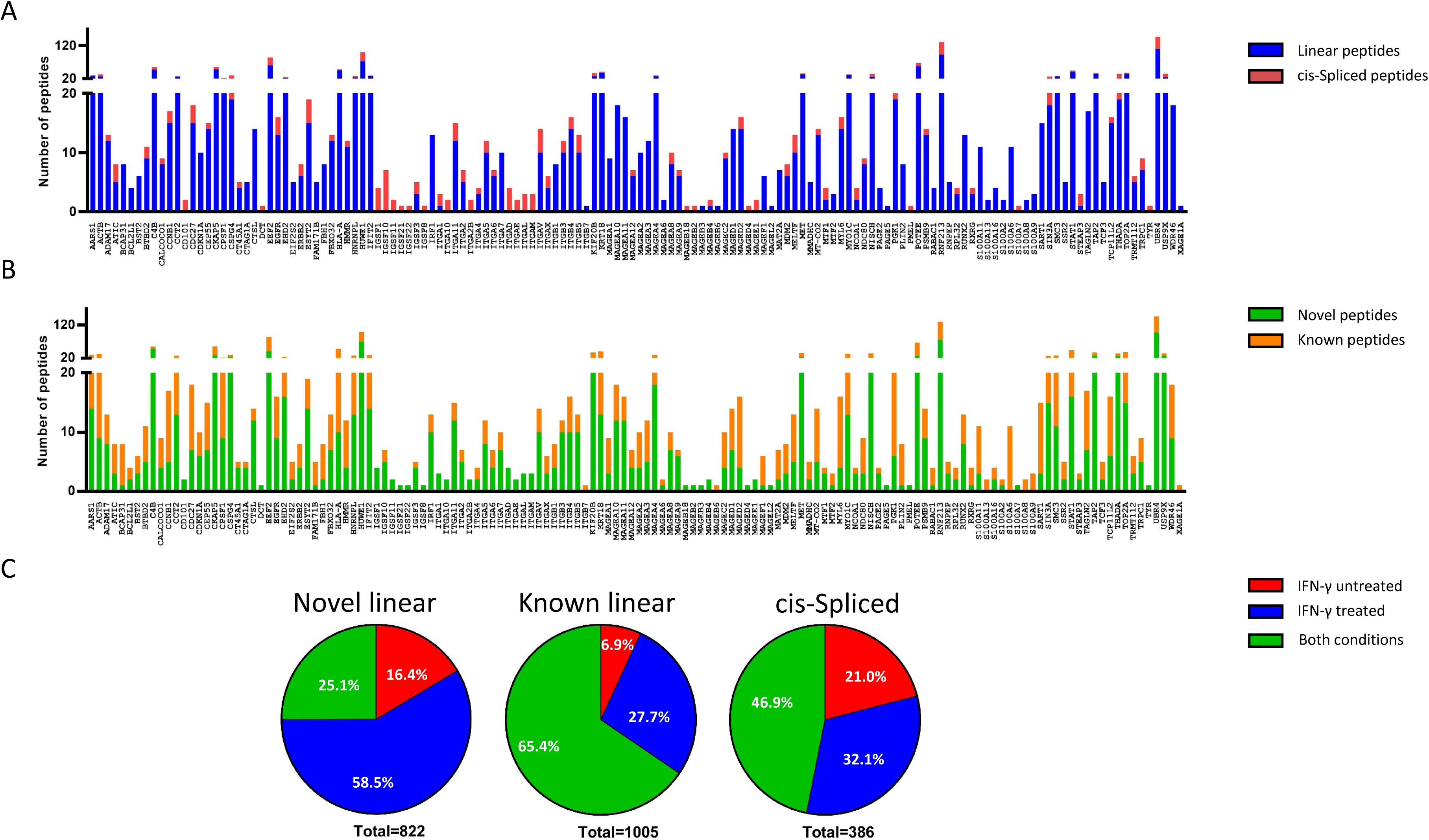
**Novel and previously described linear and spliced peptides from melanoma specific antigens were identified. A**, Both linear and *cis*-spliced peptides derived from MAA were identified in the LM-MEL-44 immunopeptidome. **B,** A substantial proportion of identified MAA have not been previously described. **C,** Proportion of MAA-associated epitopes in the presence and absence of IFN [. Almost 70% of novel identified peptides were solely present in the presence of IFNγ or generated thorough post translational splicing. IEDB website (www.iedb.or) (28), SysteMHC (https://systemhcatlas.org/)(62) and (63) were used as resources for known peptides.

### Melanoma patients expressing immunoproteasome genes have a survival advantage

Tumor recognition *in vivo* relies on the processing and generation of cognate peptides within the tumor cells. Using OncoLnc(27), we mined gene-expression data generated by the TCGA Research Network (http://cancergenome.nih.gov/) for correlation of both immuno-and constitutive-proteasome-specific genes with survival in melanoma patients. We found that expression of all three immunoproteasome-specific subunits was significantly associated with survival in melanoma patients. Conversely, constitutive proteasome-specific subunits were associated with decreased melanoma patient survival (Fig. S2).

Since immunoproteasome subunits are also expressed by immune cells, including intra-tumoral T cells that are themselves associated with better prognosis, we removed the top quartile of samples with the highest CD3 expression. Following removal of these samples, we found that a significant survival benefit, associated with 2/3 immunoproteasome subunits, was maintained (Fig. 5). Furthermore, presence of the 3 constitutive proteasome subunits was associated with a trend towards decreased survival. This indicates that patients whose tumors express an immunoproteasome have survival benefit which is specifically associated with this proteasome type. This effect persists, albeit to a lesser extent, when we removed to tumors that showed the highest CD45 infiltration, thus including non T cell lineage immune cells and APCs (Fig. 5). As immunoproteasome expression in tumors is largely driven by cytokine exposure it remains unclear if this is merely a footprint of a (previous) successful immune recognition or if it is part of the pre-conditions to allow for such an immune response.

**Figure 5.**
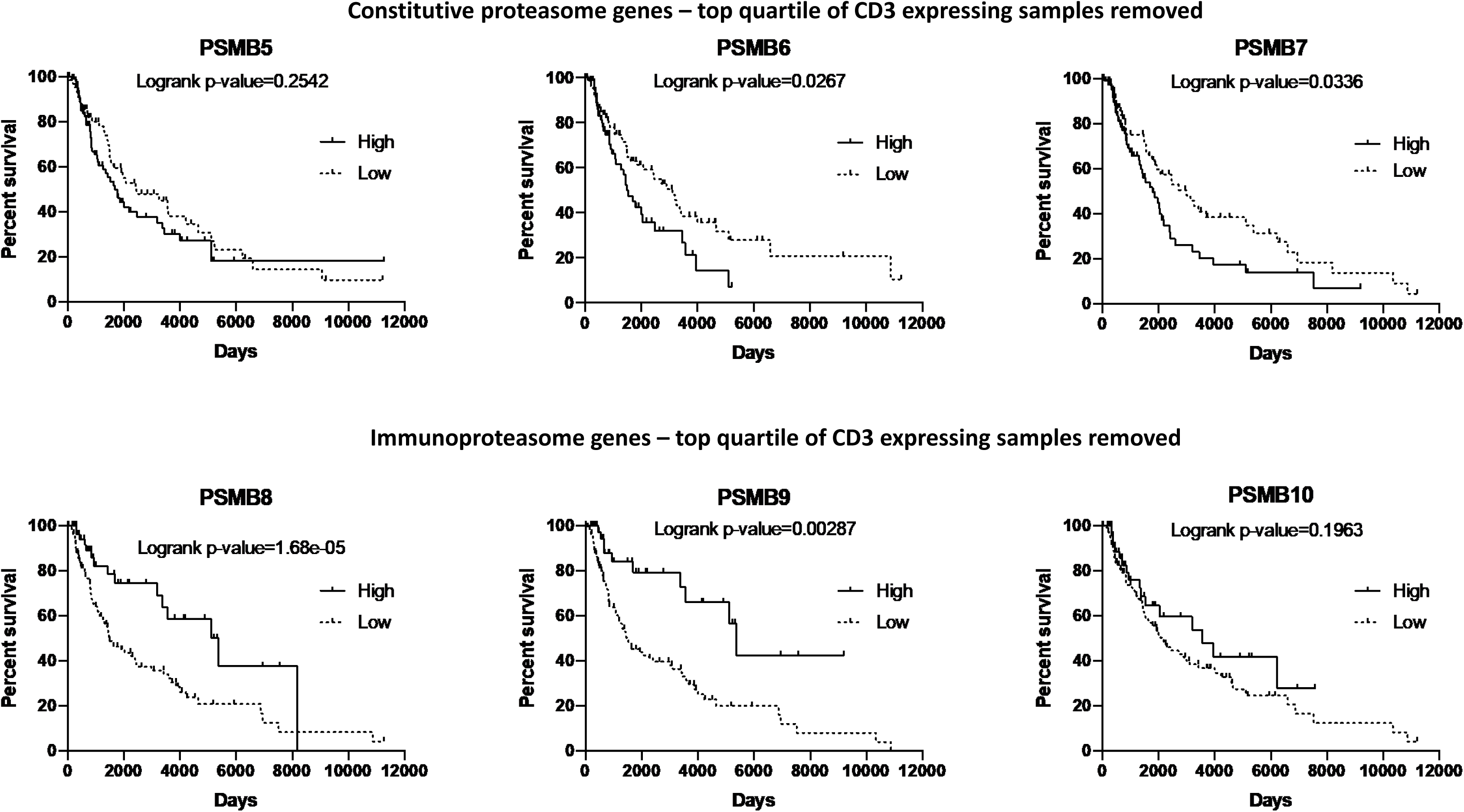
**Immunoproteasome expression is associated with survival benefit in melanoma patients.** Using the TCGA-SKCM and FM-AD datasets looking at nevi and melanomas, we selected patients whose tumors had highest (top quartile) or lowest (bottom quartile) expression of each immunoproteasome (IP) or constitutive proteasome (cP) subunits as indicated in the figure. To correct for immune infiltration, the top quartile of CD3g expressing samples were removed. The remainder of samples with high or low expression of a given proteasome subunit were plotted on the basis of patient survival over time using a Kaplan Meier survival curve (astatsa.com/LogRankTest/).

### CD8^+^ T lymphocytes frequently recognized novel linear melanoma-specific epitopes

In this study we identified several novel linear melanoma-specific peptides predicted to be bind the HLA-allotypes presented by a tumor-derived melanoma cell line (HLA-A*02:01, B*40:01/*44:02, C*03:04/*05:01). Importantly, this included HLA-A*02:01, one of the most prevalent HLA-types, and therefore a common target for peptide identification and therapeutic focus. We addressed functional immunogenicity of a selection of these peptides, by using them to stimulate CD8^+^ T lymphocytes in PBMC derived from healthy donors or melanoma patients (Fig.6, Table S5). In doing these studies, selected donors were matched for at least two HLA-allotypes, (across HLA-A/B/C) and 3 melanoma patients were matched across all three. Of note, the LM-MEL-44 cell line was derived from melanoma patient 2 and, melanoma patient 6 shares none of the HLA alleles from this cell line, serving as a negative control. Alongside these assays using melanoma antigen-derived peptides, we also assessed differences in functional immunogenicity of immuno- or constitutive-proteasome processed epitopes. This was done by pooling a selection of those peptides which were most strongly up- or down-regulated following IFN□ treatment (Fig. 6A and B, Table S5). We found that though also observed in healthy donors, CD8^+^ T lymphocyte responses to the tested peptides were more frequently seen in melanoma patients (Fig. 6A (individual) and 6B (combined donors)). Novel peptides derived from 15 of the melanoma antigens identified in our screen stimulated specific CD8^+^ T lymphocyte responses (over 2% TNFα^+^ cells) in three or more donors, demonstrating novel, functional, melanoma T cell epitopes (Fig. 6A, Table S5, S6, representative examples, Fig. S3A). Of those where a clear HLA-binding prediction could be determined, 54.5% (n=6) were predicted to bind to HLA-A*02:01, 36.4% (n=4) to HLA-B*44:02, and 9.1% (n=1) to HLA-C*05:01. The strongest responses were induced by the peptides derived from SART1 (U4/U6.U5 tri-snRNP-associated protein 1) and PGK1 (Phosphoglycerate kinase 1), both of which stimulated responses in 2-7% of T lymphocytes from 4 melanoma patients. Both of these peptides were predicted to bind to HLA-A*02:01. When a selection of peptides were pooled in groups of those up/down regulated or unchanged following IFN□ treatment, no appreciable difference in functional immunogenicity in melanoma patients was observed between groups. One pool in each group was made on the basis of higher *in silico*-predicted immunogenicity (www.iedb.org(28), Fig. 6A,B, asterisks). However, these groups did not display enhanced ability to activate CD8^+^ T lymphocytes in either melanoma patients or healthy donors.

**Figure 6:**
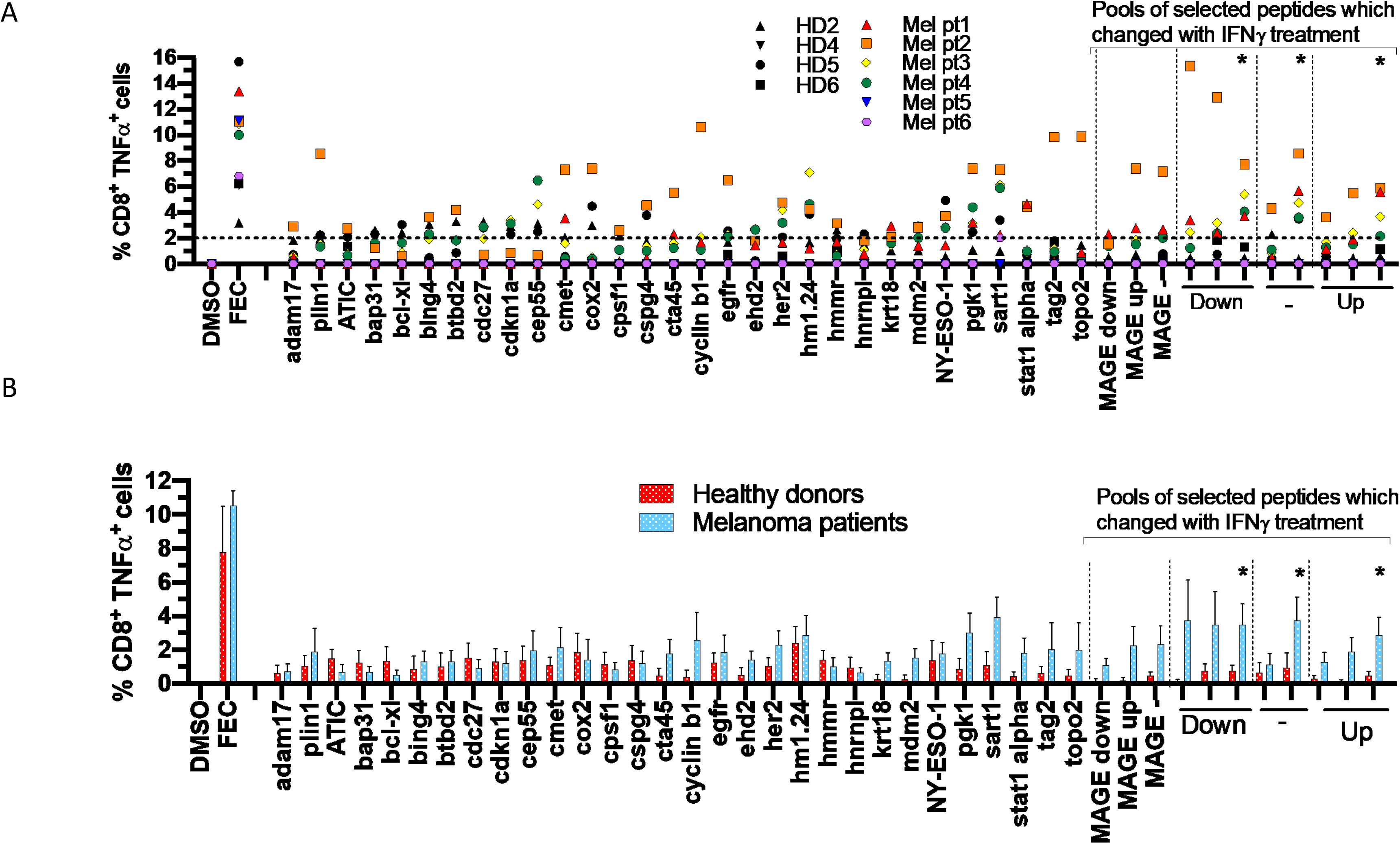
**Immunogenicity of identified melanoma-associated epitopes.** Selected MAA peptides were pooled and incubated at 10 µM final concentration with PBMC from healthy donors (n=4) or melanoma patients (n=6) for 10 days in presence of IL-2. Melanoma patient 2 is the patient from which the LM-MEL-44 cell line was derived. Melanoma patient 6 was a HLA-unmatched negative control. All other donors were HLA-matched over at least two allotypes. On day 10, CD8^+^ T lymphocytes were re-stimulated with individual peptides. Additionally cells were stimulated with pools of selected peptides whose HLA-presentation was found to be up/down regulated or unchanged in presence of IFN□ as indicated. TNFα expression measured by ICS. Data show the percentage of TNFα ^+^ CD8^+^ T lymphocytes in response to each peptide as individual values (A) or combined (B). * denotes peptides with the highest *in silico* predicted immunogenicity. In (A) all responses shown were significantly greater (p<0.05) than their respective matched DMSO control and the line denotes a conservative, arbitrary, cut-off for peptides inducing an immune response. Peptide pools outline in Table S5.

### Spliced peptides are immunogenic across patients and represent novel targets for immunotherapy

The potential implications of the presence of *s*pliced peptides for all facets of immunity have sparked intense discussions in the last 4 years(14,29–31). In cancer, their presence would dramatically widen the repertoire of potentially targetable epitopes and may allow for many more tumor-specific antigens (including mutational derived neoantigens) being presented in various HLA-contexts(16). So far only 6 immunogenic *cis*-spliced HLA-I bound peptides derived from 4 different proteins(29) have been described and most of them have been discovered by T cell assays rather than by mass spectrometry(32–37). To test some of the identified spliced peptides for their ability to activate CD8^+^ T cells *in-vitro* we synthesized 26 *cis*-spliced peptides based on (i) their *de novo* sequencing confidence score, (ii) their binding prediction score for HLA alleles expressed on LM-MEL-44 cells (HLA-A*02:01, HLA-B*40:01, HLA-B*44:02 or HLA-C*05:01) and (iii) the quality of their peptide spectrum matches (PSMs). When employed as pools of 8-9 individual peptides, all 3 pools evoked immune-responses as measured by intracellular TNFα production in CD8^+^ T cells (Fig. 7A and peptide sequences listed in Fig. 7B) in multiple melanoma patient and healthy donor derived PBMCs (example shown in Fig. S3B).

**Figure 7.**
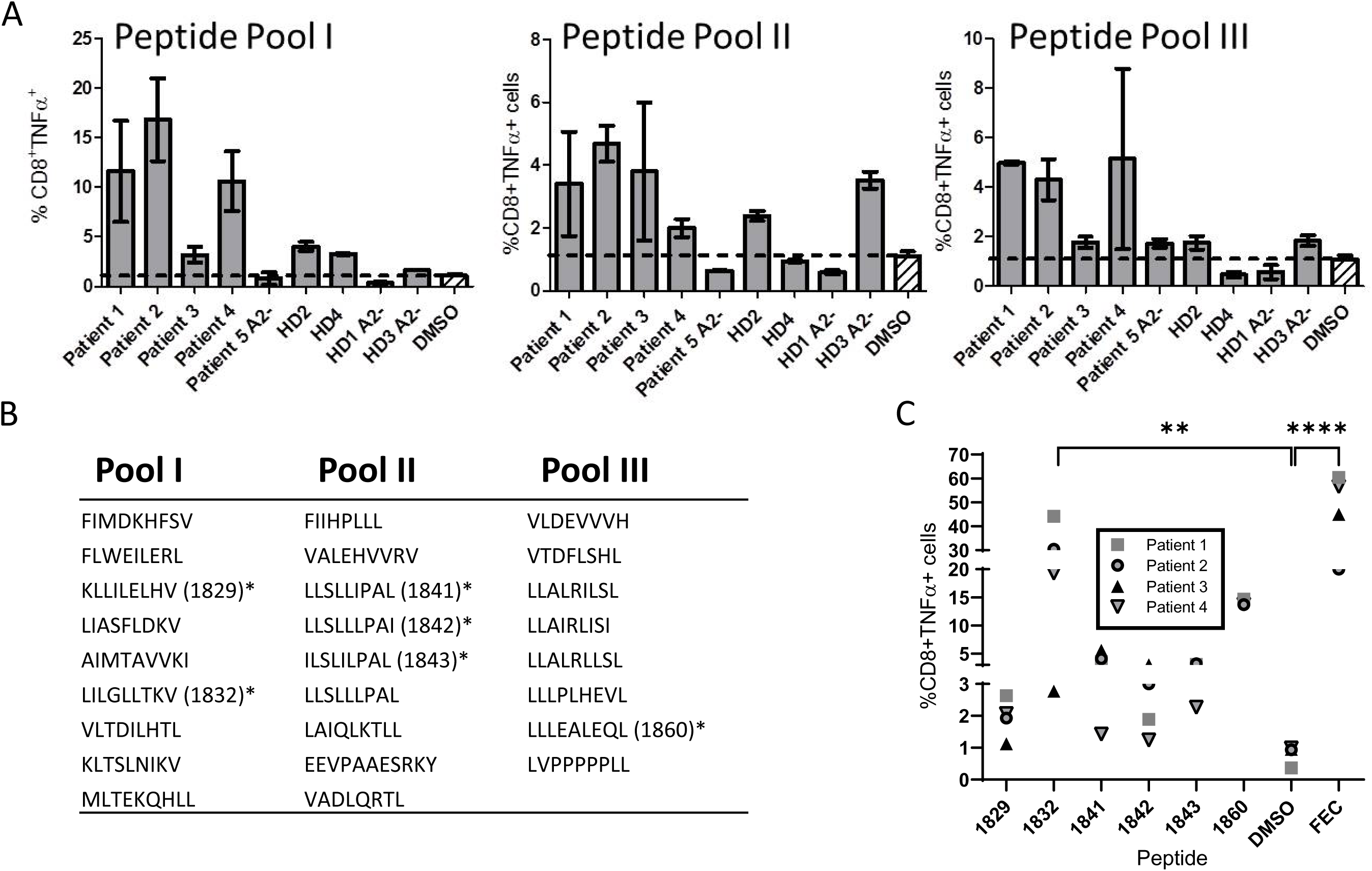
**Immunogenicity of *cis*-spliced peptide pools. A,** PBMCs from melanoma patients and healthy donors (HD) were stimulated with pooled peptides (n=8-9) for 10 days in the presence of IL-2. Cells were re-stimulated with the same pools for 8 h in the presence of BFA and TNFα expression measured by ICS. DMSO and FEC served as negative or positive control respectively. **B,** Amino acid sequence of peptide pools used in A. **C,** PBMCs from melanoma patients pre-stimulated with the pooled peptides as in A were re-stimulated with single peptides from the same pool after 12 days for 8 h in the presence of BFA and TNFα expression measured by ICS. DMSO and FEC served as negative or positive control respectively. % of TNFα ^+^ CD8^+^ cells of peptides that gave a signal above background (DMSO) are shown. Statistics was performed using Graphpad Prism as described in Material and Methods, **** p<0.0001 and ** p<0.001 after Dunnett’s multiple comparison test.

Given the differences in the potential to stimulate HLA-A2 positive vs. negative patient and healthy donor samples, most of the immunogenic peptides derived from Pools I and III in our assays seem to be HLA-A2 associated. To identify specific immunogenic peptides, PBMCs were stimulated with the listed peptide pools (Fig. 7B) for 10-12 days followed by single peptide re-stimulation. Six out of 26 peptides induced a TNF-α response above DMSO background (Fig. 7C) in more than one patient sample. Of note, the peptide demonstrating the highest immunogenicity (1832, shown as example in Fig. S3C) based on these assays is a spliced peptide derived from the cancer-testis antigen MAGE-C2 (LILGLLTKV) and showed CD8^+^ T cell activation across all 4 patients. Matched mixed effect analysis showed significant differences across peptides and peptide 1832 and FEC represented the treatments with significant differences to DMSO. However, the other shown peptides displayed higher immunogenic potential when compared to their respective DMSO control, but with very high patient-to-patient variability, as expected in these types of data, Interestingly, peptide 1832 was identified across all replicates of LM-MEL-44, -33 and -53 (Table S6). All spliced peptides that tested positive for immunogenicity in our assays were subjected to T2 peptide binding assays to examine HLA-A2 binding. As shown in Fig. S4, these peptides all stabilize HLA-A2, albeit to a lesser extent than the well described modified ELAGIGILTV HLA-A2 peptide (aa26-35) from the melanoma antigen Melan-A(38) with some just showing minor stabilization.

Of note, we did not find any particular pattern in the length of the *N*- and *C*-terminal segments of these spliced peptides nor in the distance between these segments on the protein level (Fig. S5). Taken together, these data show that these spliced peptides can serve as *bona fide* anti-cancer targets and provide a large number of additional targets would have not been considered using previous MS-based epitope discovery strategies.

## Discussion

In this study we have described a detailed and in-depth immunopeptidome presented on a patient-derived melanoma cell line (LM-MEL-44) generated from a lymph node metastasis. Our qualitative assessment of the immunopeptidome yielded around 60,000 high confidence peptide identifications that encompassed two culture conditions (+/- IFNγ) to gain insights into the influence of differences in the microenvironment of the cells on the global immunopeptidome. Furthermore, we demonstrated consistence of over 50% of these peptides with a temporally distinct autologous tumor sample, and 37% with a tumor from a different donor. The well-described effect of IFN□ in mediating changes to the composition of the antigen processing machinery, coupled with reports of differences in antigen processing between the constitutive and the immunoproteasome, led us to expect a degree of difference between the two immunopeptidomes. Nevertheless, our observation that ∼55% of linear and 47% of spliced HLA class I epitopes were exclusive to either IFNγ treated or untreated conditions, was striking. Our observations are also consistent with recent studies in ovarian and lung cancer(10, 11) that demonstrated profound changes between cytokine treatment conditions.

To have a closer look at the “tumor-specific” immunopeptidome landscape, we focussed on MAA-derived peptides. More than 50% of novel peptides that we identified were exclusively presented in the presence of IFN□. Interestingly, of the MAA epitopes that have been previously described in other studies, only 27.7% were present uniquely in IFNγ treated conditions (Fig. 4C). This observation suggests that many immunoproteasome processed epitopes may be as yet undescribed, since traditional approaches to identify tumor associated antigens have largely been undertaken using cells lines under steady state conditions (*i.e.* which express only the constitutive form of the proteasome). It is evident from our study that the steady state immunopeptidome may vary dramatically from the *in vivo* tumor scenario depending on the tumor microenvironment at any given time. Though our functional studies did not reveal a difference in the immunogenicity of peptides derived from either IFNγ treated or untreated conditions, in the *in vivo* setting a T cell response to IFNγ related epitopes is likely to be aided by correlative IFNγ influences, such as upregulation of surface HLA(39). The potential for tumor escape from CD8^+^ T lymphocyte killing due to whole scale change to the immunopeptidome upon initiation of an anti-tumor responses, and corresponding induction of IFNγ, is clear from our studies. These data become particularly significant in the context of recent studies demonstrating that tumors with an IFN□ -inflamed, or ‘hot’ microenvironment are associated with better prognosis, and are more likely to be amenable to treatment with immune checkpoint inhibitors(40). It seems conceivable that *in vivo* the difference between immunopeptidomes is indeed of immunological relevance to disease progression and overall patient prognosis. Taken together, it is tempting to speculate that antigens processed *via* the immunoproteasome may represent an untapped resource of “IFNγ-associated neo-epitopes”.

This remarkable plasticity in the peptide landscape of melanoma is further increased by the presence of spliced peptides. The identification of spliced peptides as tumor antigens in cancer was first described in 2004 in both the FGF5 protein in renal cancer(32) and the gp100 protein in melanoma(33), and since then only a further 4 *cis*-spliced peptides have been described in cancer(29). Of these 6 spliced peptides, 3 have been shown to be processed exclusively by the constitutive proteasome, and 2 by both the constitutive and immunoproteasome (and 1 undetermined)(29).

Several bioinformatics tools are now available to reliably identify spliced peptides(13,14,17,18,41). Nevertheless, the contribution of spliced peptides to the overall immunopeptidome has been reported in a range from 2 to 40% and is still heavily debated. Recent studies have identified *cis*-spliced peptides in the cancer and viral infection context (16,17,41), but few have provided experimental evidence of their immunogenicity (41). In this study we identified 386 *cis-*spliced peptides that were potentially derived from MAA and therefore considered as potential candidates to induce therapeutic immune responses. We demonstrated that generation and presentation of spliced peptides is not a random process since within three biological replicates we found comparable reproducibility of both linear and spliced peptides. Interestingly, more than 50% of spliced peptides identified in the LM-MEL-44 cell line were present on at least one of the other two distinct cell lines (LM-MEL-53 and 33). In addition, we demonstrated the immunogenicity of 6 *cis*-spliced epitopes tested across multiple patients, strengthening the argument that *cis*-splicing is a random, functional process leading to diversification of the antigenic pool of peptides. In how far increased potential to evoke CD8+ T cell activation reflects meaningful and translatable anti-tumour effects remains to be tested in much larger patient cohorts with additional clinical data and in a more formal setting.

This study and other recent publications which focused on the identification of spliced peptides (16,42,43) and their impact on the plasticity of the immunopeptidome, will beyond doubt open up new questions and opportunities in the field. These will range from the basic understanding of immune-tolerance, autoimmunity and thymic selection to opportunities for development of novel peptide-based therapeutics. This includes vaccines in an infectious and cancer setting where predictability and HLA-binding characteristics of linear and constitutive proteasome-derived peptides were potentially limiting factors.

## Materials and Methods

### Human ethics approval

Samples used in this study were derived from patients who consented to participate in a clinical research protocol approved by Austin Health Human Research Ethics Committee (HREC H2006/02633).

### Melanoma cell line culture

Establishment and characterization of the melanoma cell lines used has been previously described(44, 45). Cells were cultured in RF10 consisting of RPMI 1640, 2 mM Glutamax, 100 IU/ml Penicillin, 100 µg/ml Streptomycin and 10% heat-inactivated fetal calf serum (all Invitrogen). For induction of immunoproteasome catalytic subunits, cells were incubated with 100 ng/ml IFNγ (Peprotech) for 72 h prior to experiments.

### Melanoma cell line sequencing

Whole exome sequencing of the LM-MEL-44 cell line was performed using the NimbleGen EZ Exome Library v2.0 kit and run on a Illumina Hiseq2000 instrument as previously described(46). Sequence reads were aligned to the human genome (hg19 assembly) using the Burrows–Wheeler Aligner (BWA) program(47). Single nucleotide variants (SNVs) and indels were identified using the GATK Unified Genotyper(48), Somatic Indel Detector(49) and MuTect (Broad Institute)(50).

### Isolation of peptides bound to HLA class I molecules

HLA class I peptides were eluted from LM-MEL-44, 33 and 53 cells (prior to or after treatment with IFNγ) as described previously(51–54). In brief, for replicate one of LM-MEL-44, 3 x 10^9^ cells were lyzed in 0.5 % IGEPAL, 50 mM Tris-HCl pH 8.0, 150 mM NaCl supplemented with protease inhibitors (CompleteProtease Inhibitor Cocktail Tablet; Roche Molecular Biochemicals) for 45 min at 4 °C. Lysates were cleared by ultracentrifugation at 40,000 *g* and HLA class I complexes were immunoaffinity purified using DT9 (anti HLA-C) and W6/32 (pan anti-HLA-I) monoclonal antibodies. For replicate two and three of LM-MEL-44 and also LM-MEL-33 and LM-MEL-53, 5 x 10^8^ cells (for each sample) were lyzed in 0.5 % IGEPAL, 50 mM Tris-HCl pH 8.0, 150 mM NaCl supplemented with protease inhibitors for 45 min at 4 °C. Lysates were cleared by ultracentrifugation at 40,000 *g* and HLA class I complexes were immunoaffinity purified using W6/32 (pan anti-HLA-I) monoclonal antibody.

### Fractionation of HLA-bound peptides by reversed-phase high-performance liquid chromatography (RP-HPLC)

The HLA-peptide eluates were loaded onto a 4.6 mm internal diameter x 50 mm monolithic C18 RP-HPLC column (Chromolith Speed Rod; Merck) at a flow rate of 1 ml/min using an EttanLC HPLC system (GE Healthcare) with buffer A (0.1 % trifluoroacetic acid (TFA)) and buffer B (80 % ACN / 0.1 % TFA) as mobile phases. The bound peptides were separated from the class I heavy chains and ß2m molecules using an increasing concentration of buffer B. Peptide-containing fractions (500 µl) were collected, vacuum concentrated to ∼5 µl and combined into nine pools, reconstituted to 12 µl with 0.1 % formic acid (FA). Indexed retention time (iRT) peptides(55) were spiked in for retention time alignment.

### Identification of HLA bound-peptides using data-dependent acquisition (DDA)

For the first replicate of LM-MEL-44 we used a Dionex UltiMate 3000 RSLCnano system equipped with a Dionex UltiMate 3000 RS autosampler, the samples were loaded via an Acclaim PepMap 100 trap column (100 µm x 2 cm, nanoViper, C18, 5 µm, 100 Å; Thermo Scientific) onto an Acclaim PepMap RSLC analytical column (75 µm x 50 cm, nanoViper, C18, 2 µm, 100 Å; Thermo Scientific). The peptides were separated by increasing concentrations of 80 % ACN / 0.1 % FA at a flow of 250 nl/min for 65 min and analyzed with a QExactive Plus mass spectrometer (Thermo Scientific). In each cycle, a full ms1 scan (resolution: 70.000; AGC target: 3e6; maximum IT: 120 ms; scan range: 375-1800 m/z) preceded up to 12 subsequent ms2 scans (resolution: 17.500; AGC target: 1e5; maximum IT: 120 ms; isolation window: 1.8 m/z; scan range: 200-2000 m/z; NCE: 27). To minimize repeated sequencing of the same peptides, dynamic exclusion was set to 15 s and the ‘exclude isotopes’ option was activated.

For second and third replicates of LM-MEL-44, LM-MEL-33 and LM-MEL53 we used a Dionex, Sunnyvale, CA, UltiMate 3000 RSLCnano system equipped with a Dionex UltiMate 3000 RS auto sampler, the samples were loaded via an Acclaim PepMap 100 trap column (100 μm × 2 cm, nanoViper, C18, 5 μm, 100å; Thermo-Fisher Scientific, Waltham, MA) onto an Acclaim PepMap RSLC analytical column (75μ m × 50 cm, nanoViper, C18, 2μm, 100 Å; ThermoFisher Scientific). The peptides were separated by increasing concentrations of 80 % ACN/0.1 % FA at a flow of 250 nL/min for 158 min and analyzed with an Orbitrap Fusion™ Tribrid™ mass spectrometer (ThermoFisher Scientific). 6 μL of each sample fraction was loaded onto the trap column at a flow rate of 15 μL/min.

Orbitrap Fusion™ Tribrid™ mass spectrometer (ThermoFisher Scientific) was set to data-dependent acquisition mode with the following settings: All MS spectra (MS1) profiles were recorded from full ion scan mode 375-1800 m/z in the Orbitrap at 120,000 resolution with automatic gain control (AGC) target of 400,000 and dynamic exclusion of 15 s. The top 12 precursor ions were selected using top speed mode at a cycle time of 2 s. For MS/MS, a de*cis*ion tree was made which helped in selecting peptides of charge state 1 and 2-6 separately. For single charged analytes only ions falling within the range of m/z 800-1800 were selected. For +2 to +6 m/z s no such parameter was set. The c-trap was loaded with a target of 200,000 ions with an accumulation time of 120 ms and isolation width of 1.2 amu. Normalized collision energy was set to 32 (high energy collisional dissociation (HCD)) and fragments were analyzed in the Orbitrap at 30,000 resolution.

### DDA data analysis

Linear and *cis*-spliced peptide sequences were identified as described previously(14). In brief, the acquired .raw files from six LM-MEL-44 line were searched with PEAKSStudio X (Bioinformatics Solutions Inc., Waterloo, Ontario, Canada) against the human UniProtKB/SwissProt (v2017_10) database, which was manually corrected for the single nucleotide variants characteristic of the LM-MEL-44 cell line as identified by whole exome sequencing. The parent mass error tolerance was set to 10 ppm for *de novo* sequencing and database search and the fragment mass error tolerance to 0.02 Da for both searches. Oxidation of M and deamidation of N & Q were set as variable modifications and a FDR cutoff of 1% was applied. High confidence *de novo* peptide sequences without any linear peptide match in the provided database were further interrogated with the “Hybrid finder” algorithm and the identified *cis*-spliced candidate sequences from all 6 samples combined together and added back to the original UniProtKB/SwissProt database and all data researched using PEAKS DB. Linear and cis-spliced peptides in this search were extracted at 5% FDR to create the final list of identified peptides. For identification of linear and *cis* spliced peptides from LM-MEL-33 and 53, we used this combined database and the same setting on PEAKS studio.

### Binding prediction

We used the NetMHC4 (56, 57) algorithms for binding predictions for both spliced and linear peptides. A default rank cut-off of 2 was implemented as a binder peptide.

### Immunogenicity prediction

We used the Immune Epitope Database and Analysis Resource (IEDB: www.iedb.org), Class I Immunogenicity algorithm for immunogenicity predictions of linear peptides(58). The peptides used in functional assays indicated as “predicted immunogenic” had an immunogenicity score of between 0.37 and 0.55 (Table S5).

### T cell stimulation assay

To assess T cell responses selected peptides were synthesized (Mimotopes, VIC, Australia). PBMC from healthy donors (Australian Red Cross, VIC, Australia), or melanoma patients (Austin Health HREC approved protocol HREC H2006/02633) were purified by density centrifugation over Ficoll Hi-Paque. Cells were cultured in TCRPMI consisting of RPMI 1640, 2 mM Glutamax, 100 IU/ml Penicillin, 100 µg/ml Streptomycin, 20 mM HEPES, 1% nonessential amino acids, 1 mM sodium pyruvate, 55 µM β-mercaptoethanol, and 10% human AB serum (Australian Red Cross, VIC, Australia). Peptides were combined into pools of 5-9 peptides as outlined in Supplementary Table 6. 10^6^ PBMC/ml were incubated with 10 µM of each peptide in pools for 10 - 12 days at 37 °C. IL-2 (100 IU/ml) was added and replaced every 3 days. Statistics were performed using GraphPad Prism. For peptide responses in Figure 6a a repeated-measures 2-way ANOVA was used, and for *cis*-spliced peptide assays in Figure 7 a matched mixed effect analysis with Dunnett’s multiple comparison test was used to compare each column to DMSO.

### Intracellular cytokine staining (ICS) of antigen-activated T-lymphocytes

To assess antigen responses, T-lymphocytes were restimulated (following 10 - 12 days incubation as outlined above) with peptide pools for 4-8 h in TCRPMI in presence of 10 µg/ml brefeldin A (BFA, Golgi plug). Cells were washed with PBS (Invitrogen) labeled with live/dead fixable violet stain (Invitrogen), then incubated with antibodies against CD3 and CD8 (BD biosciences) for 15 min at 4°C. Samples were washed and fixed for 20 min at 4°C. Cells were permeabilized and stained with anti-TNFα (eBiosciences) for 25 min at 4°C. The gating strategy was: SSC/FSC; Singlets; SSC/LD^-^; CD3^+^/CD8^+^; CD8^+^/TNFα^+^(Fig. S6). Data were acquired on a FACSCanto (BD biosciences, VIC Australia) and analyzed with FlowJo software (Version 10, FlowJo, Ashland OR, USA). To account for the large variation in DMSO background CD8^+^ T cell activity across multiple donors, signals were normalized by subtracting the background from DMSO control treated samples in each case.

### HLA-A2 stabilization assays

The binding activity of the peptides was assayed by measuring peptide-induced stabilization of HLA-A2 on TAP-deficient T2 cells by flow cytometry. T2 cells were cultured in RF-10 (RPMI with 10% serum, 5% Glutamine, 5% Pen/Strep) in T25 flasks for 2 - 3 days before the assay. T2 cells (2x 10^5^ cells/well) were cultured for 16 h at 37 °C in 5 % CO_2_ in 200 µL RF10 in 96-well U-bottomed plates in presence or absence of 10 µg/ml of synthetic peptides. Peptides from Melan-A (modified aa26-35 ELAGIGILTV and aa 60-72) served as positive or negative controls respectively. All peptides were tested in triplicate.

After 16 h stimulation the cells were washed and stained with anti-HLA A2 monoclonal antibody BB7.2 (Biolegend) for 30 min. at 4 °C. Cells were subsequently stained with Fixable Viability Kit (Zombie NIR™, Biolegend) for 15 min. at 4 °C before flow cytometry on a FacsCanto (BD). Data was analyzed using FlowJo software (Version 10, FlowJo, Ashland OR, USA).

### Validation spliced peptides using retrospectively synthesized peptides and retention time prediction

We validated the identity of a panel of spliced peptides (including peptides that were tested for immunogenicity), using 28 synthetic peptides (Mimotopes, Melbourne, Australia) by comparing chromatographic retention and MS/MS spectra with the original p-HLA (Fig. S7). The PKL files of the synthetic peptide and corresponded eluted peptides were exported from PEAKS X studio software. For evaluating the similarity between two spectra, we predicted all b- and y-ions for each sequence and then extracted the corresponding intensity for each ion (with a fragment mass error tolerance of 0.1 Da). The Pearson correlation coefficient and the corresponding p-value (Prism version 8.01, GraphPad) between the log10 intensities of identified b- and y-ions in the synthetic and sample derived spectra were calculated (Fig. S7 and S8) (59). The closer a correlation coefficient to 1, the more identical the spectra. All tested peptides were found to have a *p*-value of less than 0.05.

We calculated the iRT index of each synthetic peptide and corresponding eluted peptide using the retention time of a standard set of reference peptides (iRT kit, Biognosys) that were spiked into all samples (Fig. S8) (55).

We also used GPTime tool (59) to compare the predicted versus actual chromatographic retention time of both the identified linear and spliced peptides. For each data set generated from LM-MEL-44 cell lines (6 replicates in total) we sorted peptides (8-12 mer peptides without modification) based on the –logP score (high to low) from Peaks Studio software. We used the first 1000 peptides for training the algorithm and then used the trained algorithm to predict the retention times of all linear and spliced peptides in the corresponding dataset (Fig. S9). We also calculated Grand Average of Hydropathy (GRAVY) Score of all identified 8-12 mer peptides (Table S1) (60).

## Acknowledgments

The authors acknowledge the Monash Proteomics & Metabolomics Facility for the provision of mass spectrometry instrumentation, training and technical support, as well as the Monash University Flowcore for flow cytometry instrumentation and assistance.

This project was funded in part by Ludwig Cancer Research, Melanoma Research Alliance (MRA), the Victorian Cancer Agency supported Melbourne Melanoma Project (MMP), Australian National Health and Medical Research Council (NHMRC)

Project Grants 1007381 and 1165490 (to AWP), and the Victorian State Government Operational Infrastructure Support Program. AB is supported by a grant from La Trobe University (RFA Understanding Disease). CL is supported by an Australian NHMRC CJ Martin Early Career Fellowship 1143366. JC was supported by Australian NHMRC Practitioner Fellowship 487905. AWP is supported by NHMRC Principal Research Fellowship 1137739. AB is the recipient of a Fellowship from the Victorian Government Department of Health and Human Services acting through the Victorian Cancer Agency.

Computational resources were supported by the R@CMon/Monash Node of the NeCTAR Research Cloud, an initiative of the Australian Government’s Super Science Scheme and the Education Investment Fund.

The mass spectrometry proteomics data have been deposited to the ProteomeXchange Consortium via the PRIDE (61) partner repository with the dataset identifier PXD014397 and 10.6019/PXD014397 (Username, Password: LIBn7t49).

## Supplementary Figures

**Figure S1.**
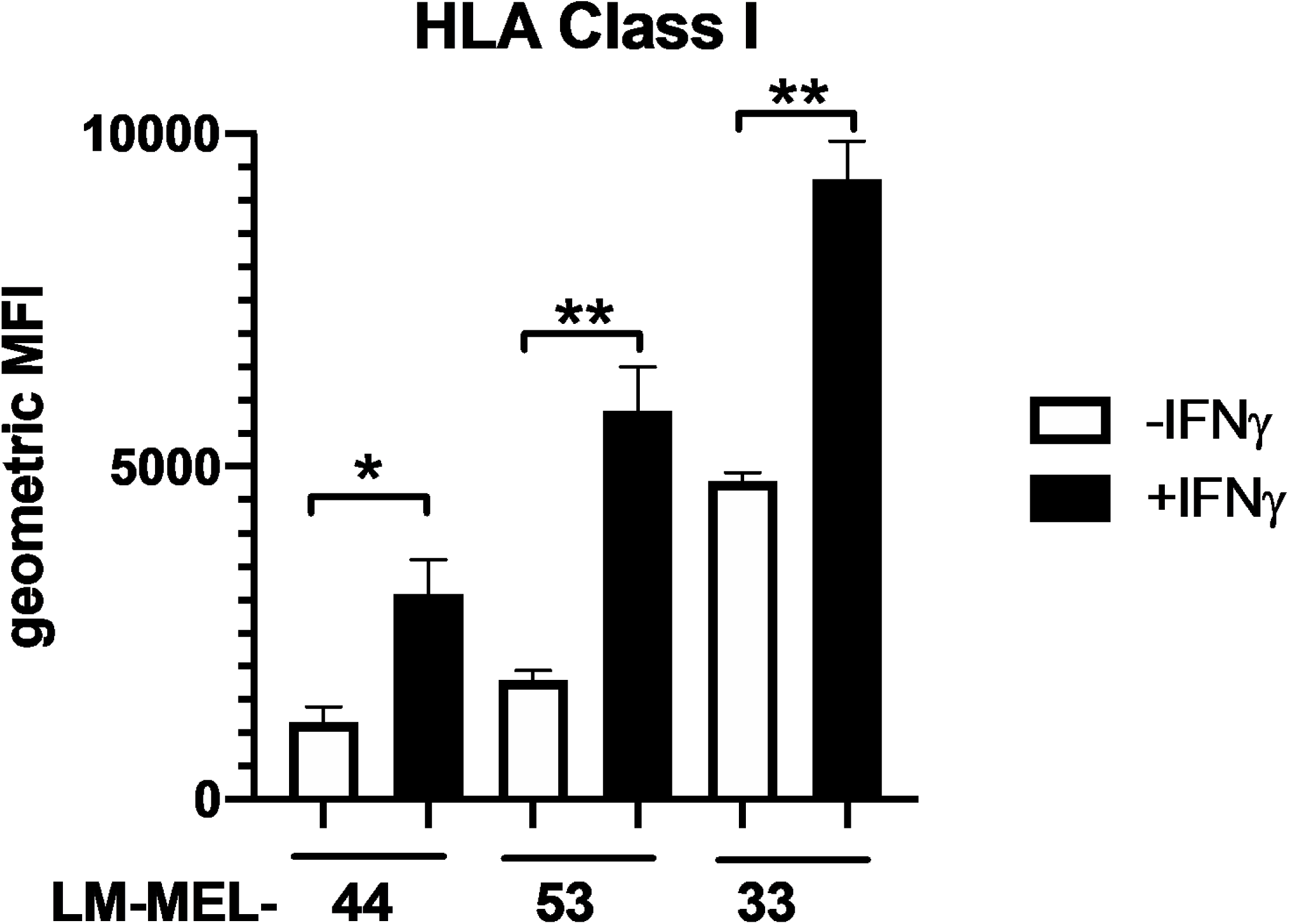
**IFN**γ **treatment increase the expression of HLA-I molecules.** The melanoma cell lines LM-MEL-33, -44, and -53 were cultured in presence/absence of 100 ng/ml IFNγ for 72 h. Cells were labelled with anti-pan HLA-I and the expression of HLA-I on the cell surface was determined by flow cytometry. *=p<0.05, **=p<0.001.

**Figure S2.**
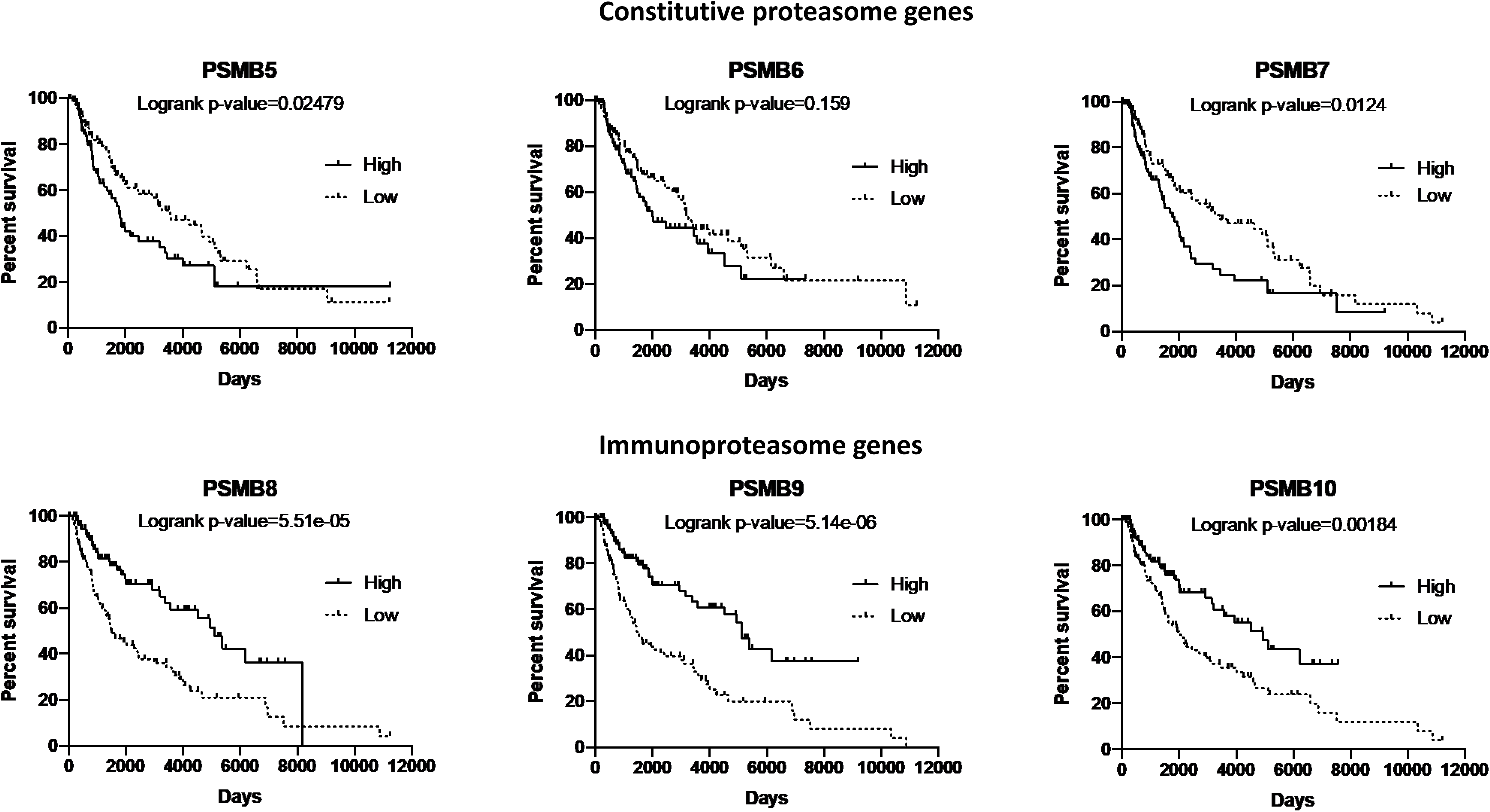
Using the TCGA-SKCM and FM-AD datasets looking at nevi and melanomas, we selected patients whose tumors had highest (top quartile) or lowest (bottom quartile) expression of each immunoproteasome (IP) or constitutive proteasome (cP) subunits as indicated in the figure. Samples with high or low expression of a given proteasome subunit were plotted on the basis of patient survival over time using a Kaplan Meier survival curve (astatsa.com/LogRankTest/).

**Figure S3.**
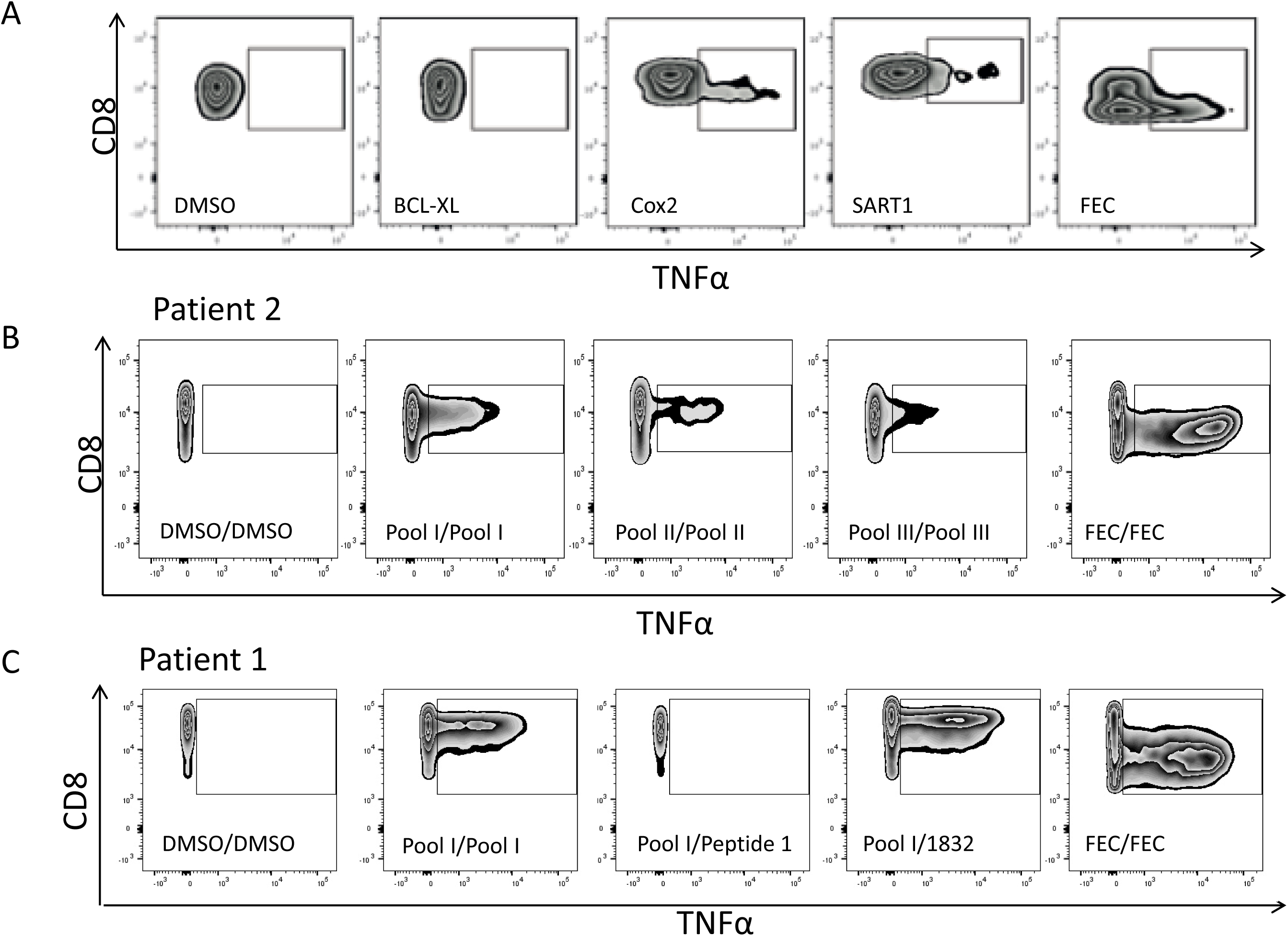
**The FACS plot examples for the peptide stimulations. A,** FACS plot showing TNFα vs CD8 in selected patients re-stimulated with either linear peptides. **B,** pools of spliced peptides. **C,** or single spliced peptides. DMSO stimulations served as negative and background control and FEC as positive control. Labelling denotes first stimulation and re-stimulation.

**Figure S4.**
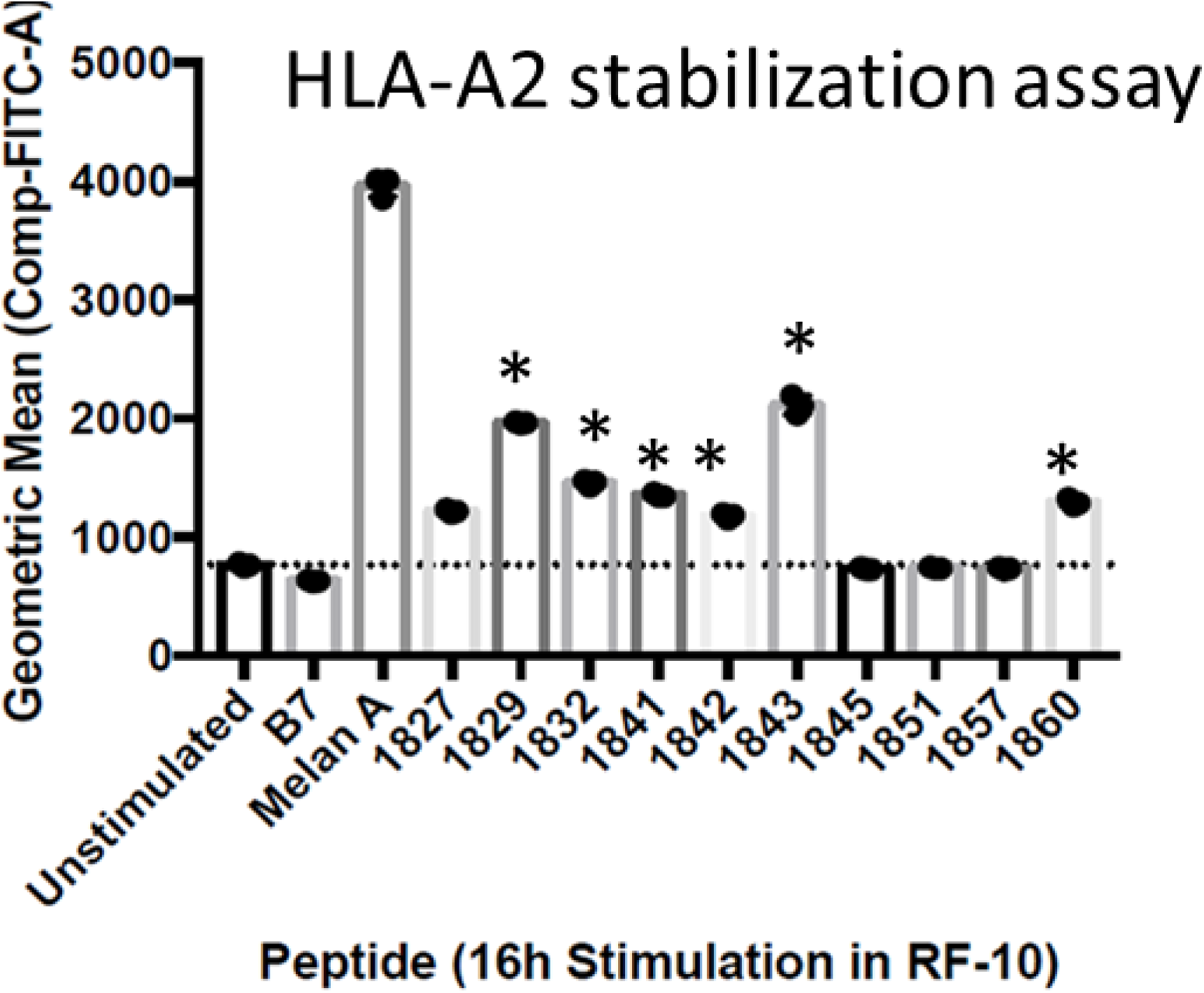
*Cis*-spliced peptides that stimulated CD8^+^ T cells as measured by TNFα to a higher degree than DMSO in at least 1 patient plus some randomly picked *cis*-spliced peptides were subjected to HLA-A2 stabilization assays on T2 cells as described in M&M. None HLA-A2 binding peptides (B7) or the Melan-A modified HLA-A2 epitope served as negative and positive control respectively.* denotes immunogenic peptides.

**Figure S5.**
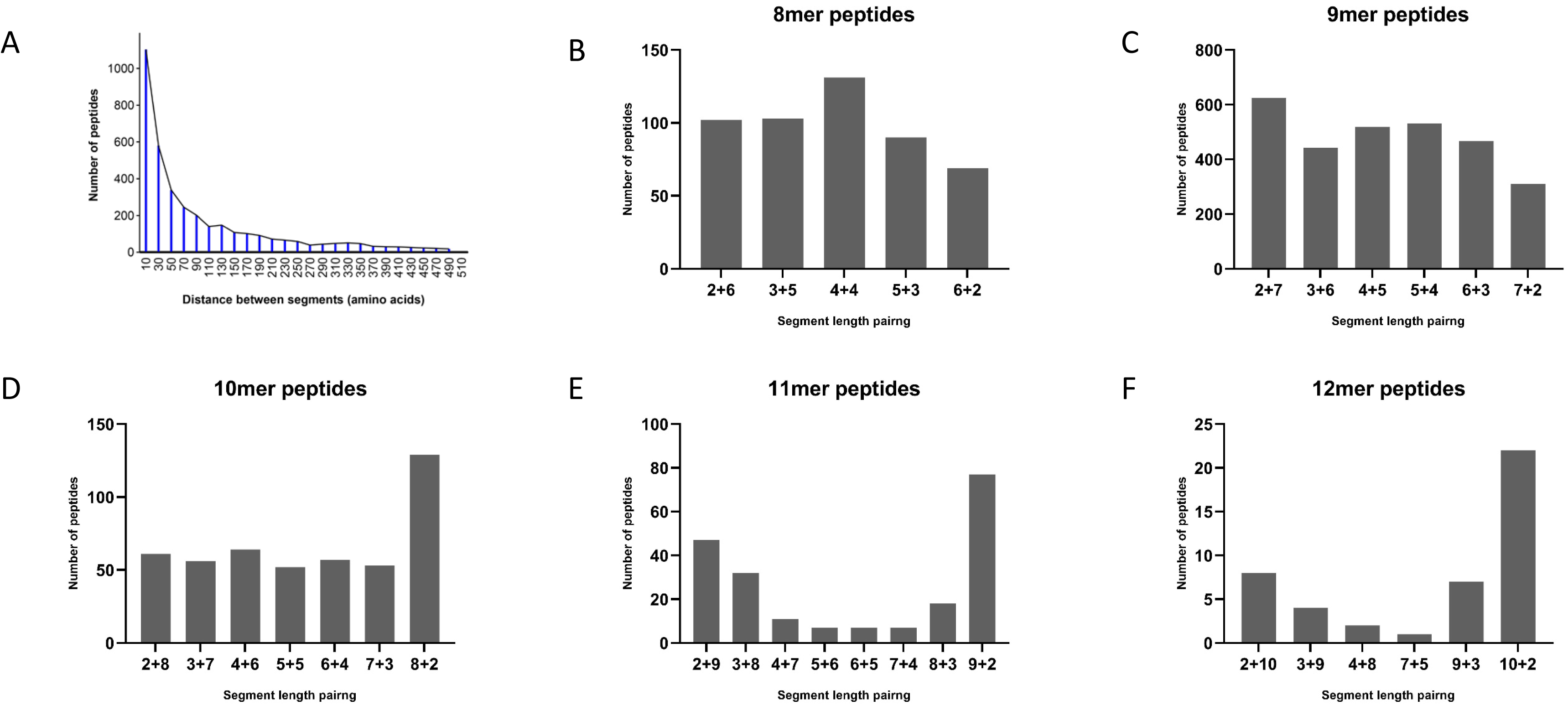
A, Length distribution of the region separating the N- and C-terminal segment of a cis-spliced peptide on the protein level (in amino acids). B-F, Length distribution of N- and C-terminal segments of *cis*-spliced peptides, shown for 8-12 mers (in amino acids).

**Figure S6.**
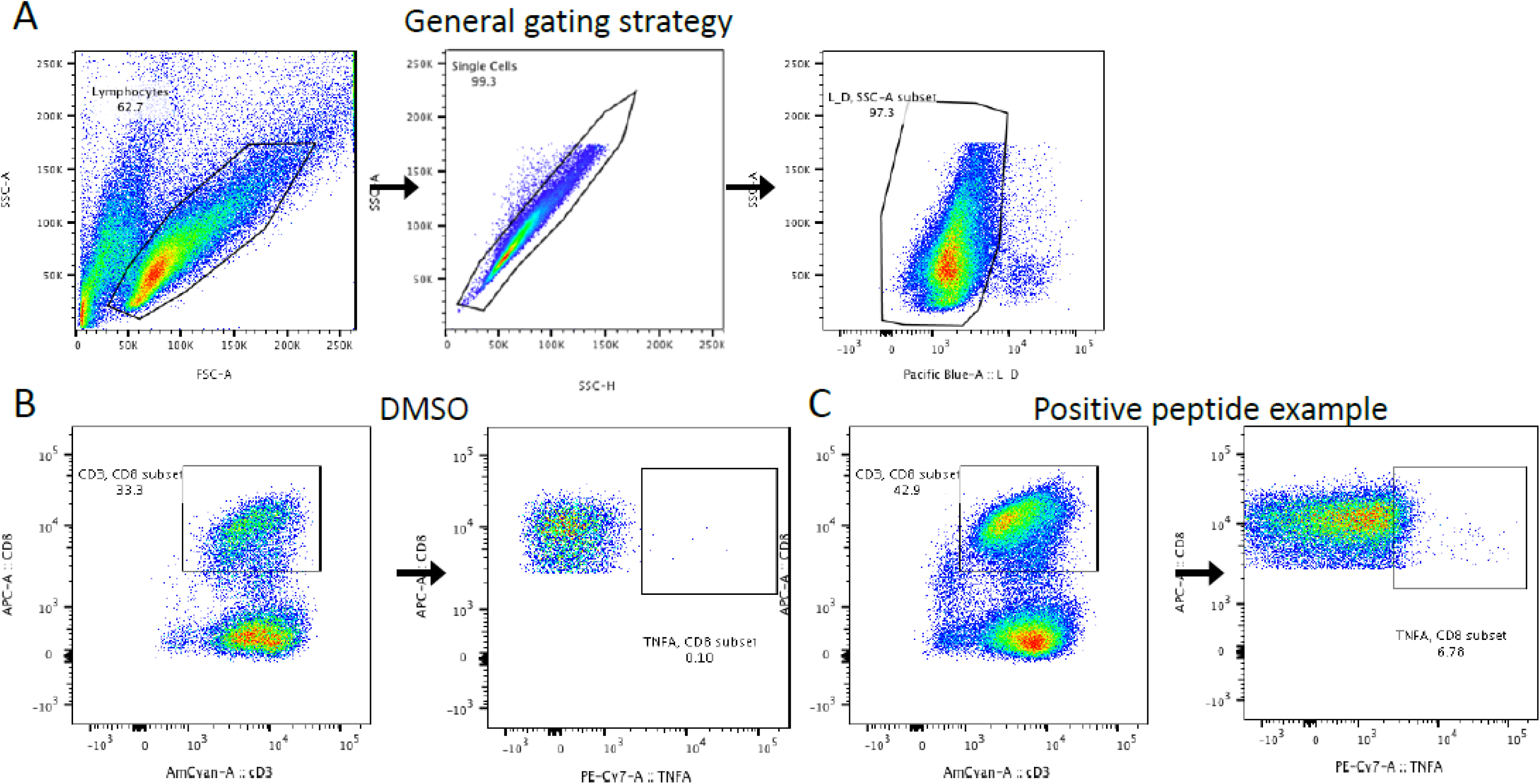
Flow cytometry gating strategy. **A**, Cells were gated based on size (FSC-A/SSC-A), doublets and dead cells were excluded, and this strategy was used for all samples. The DMSO-treated negative control. **B,** was used to set the TNF ^+^/CD8^+^ gate after CD3^+^/CD8^+^ gating. An example of a peptide-specific CD8^+^ T cell response is shown (C).

**Figure S7.**
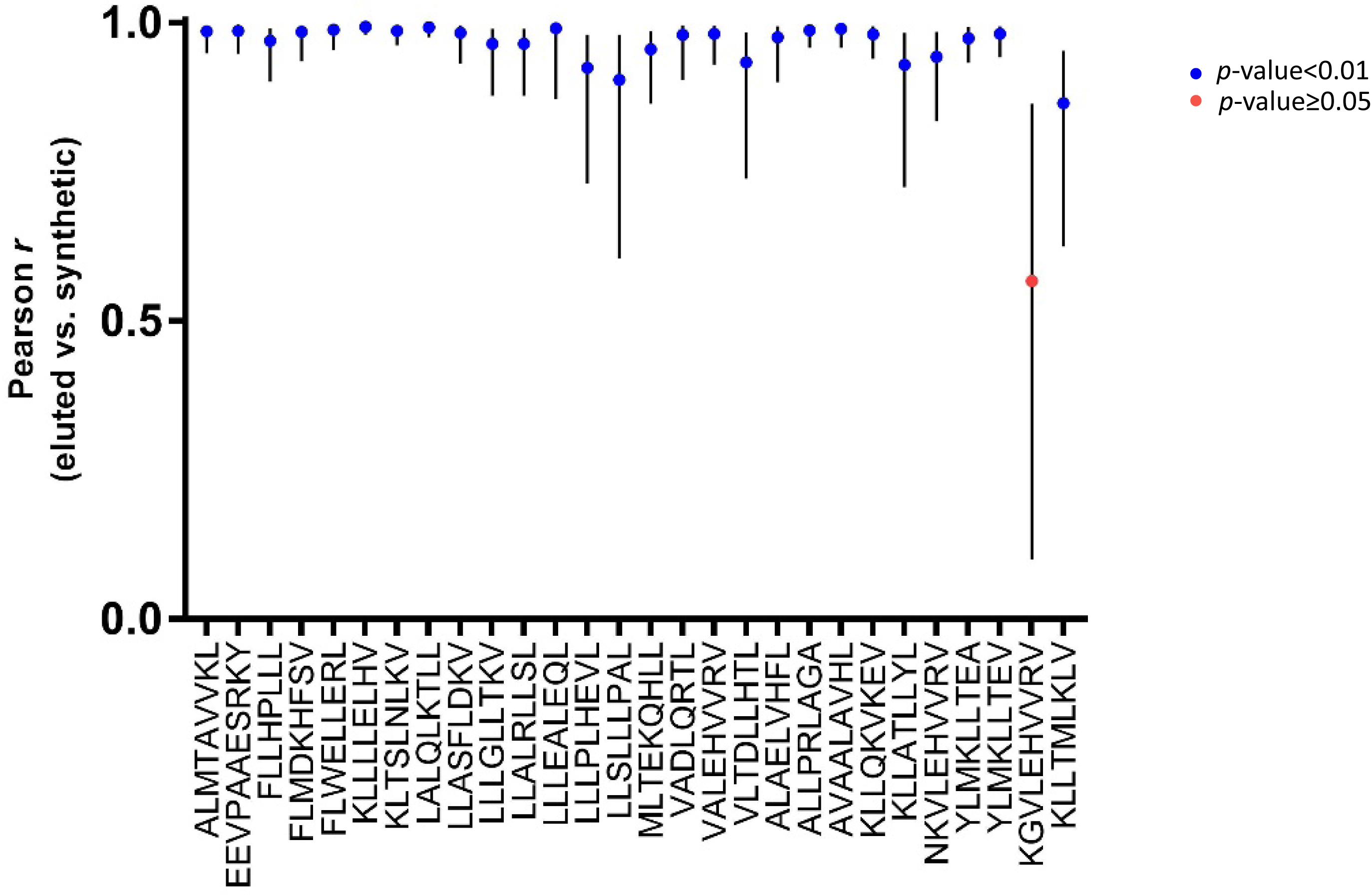
Pearson r value correlation score of b and y ions in spectra from 28 eluted *cis*-spliced peptides from LM-MEL-44 cell line compared with their synthetic versions. This analysis approved the authenticity of 27 sequences and disapproved one sequence.

**Figure S8.**
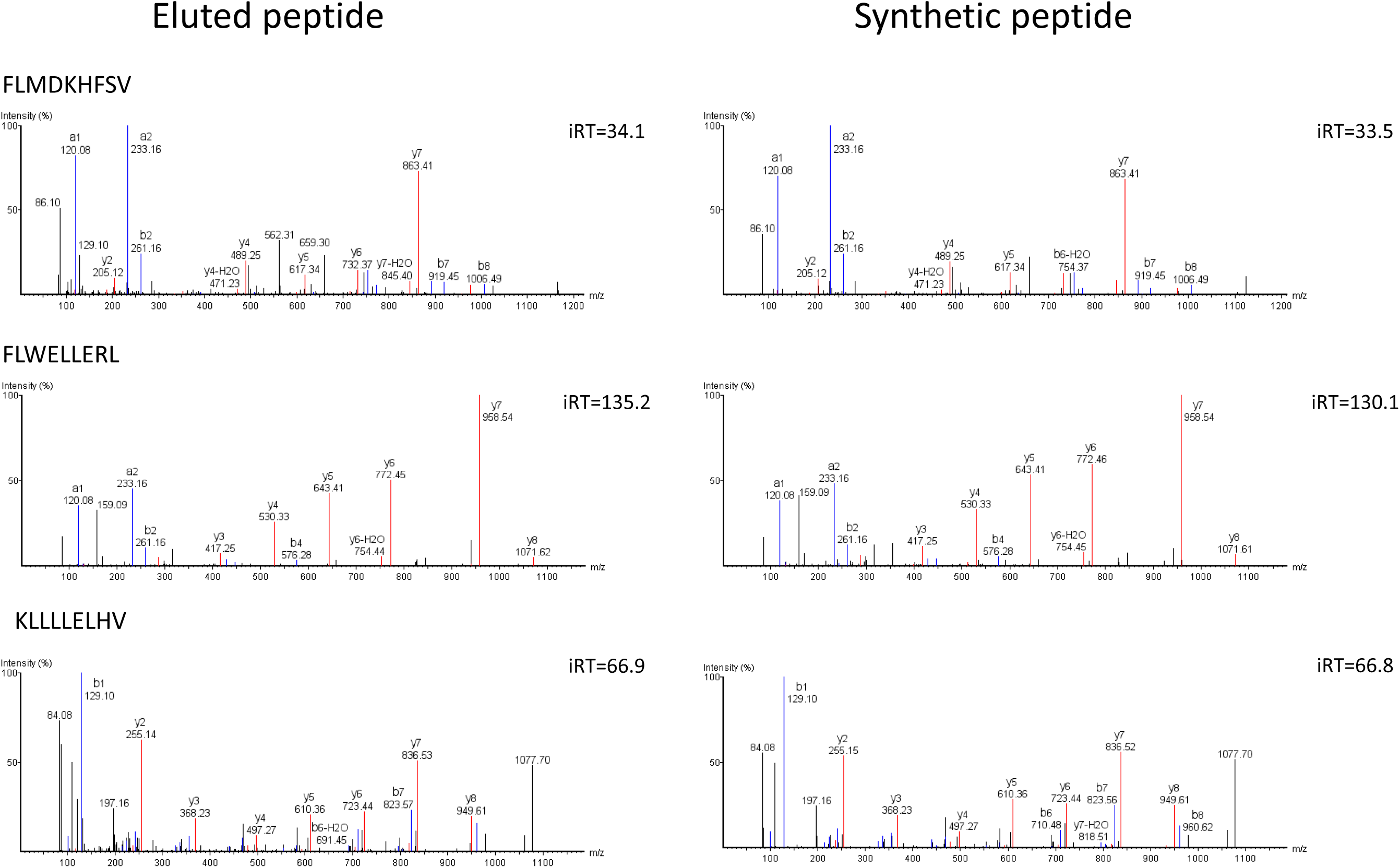
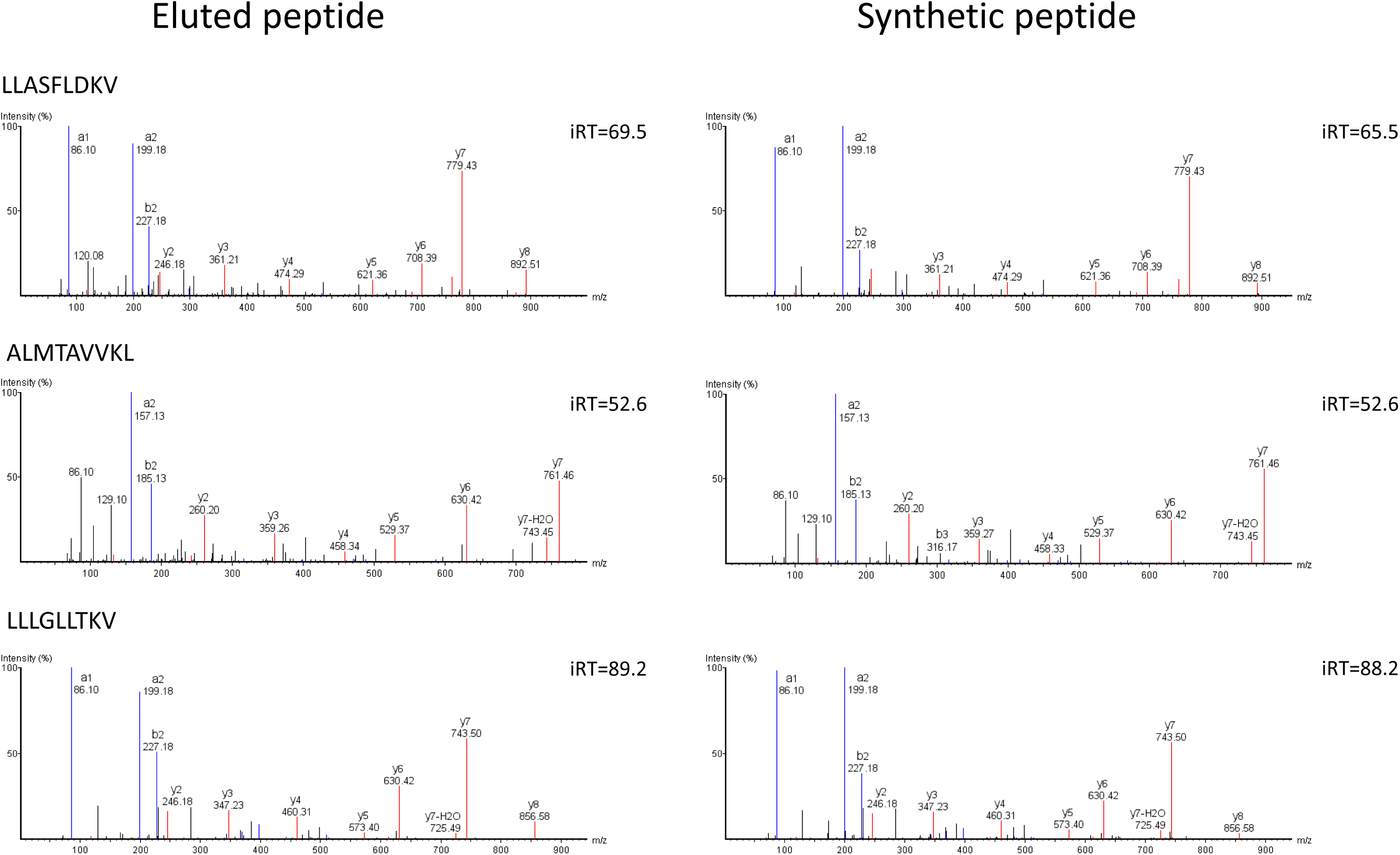
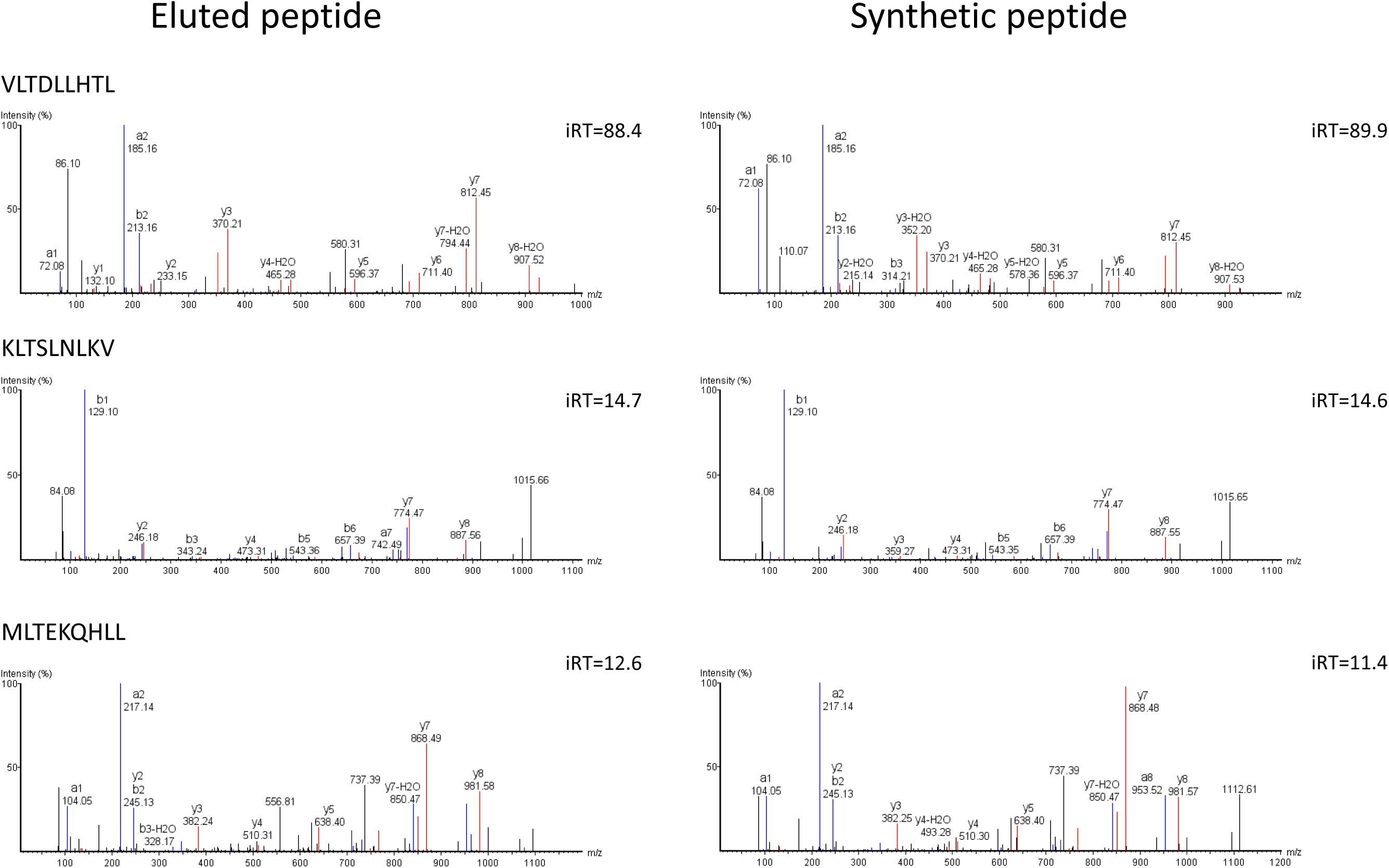
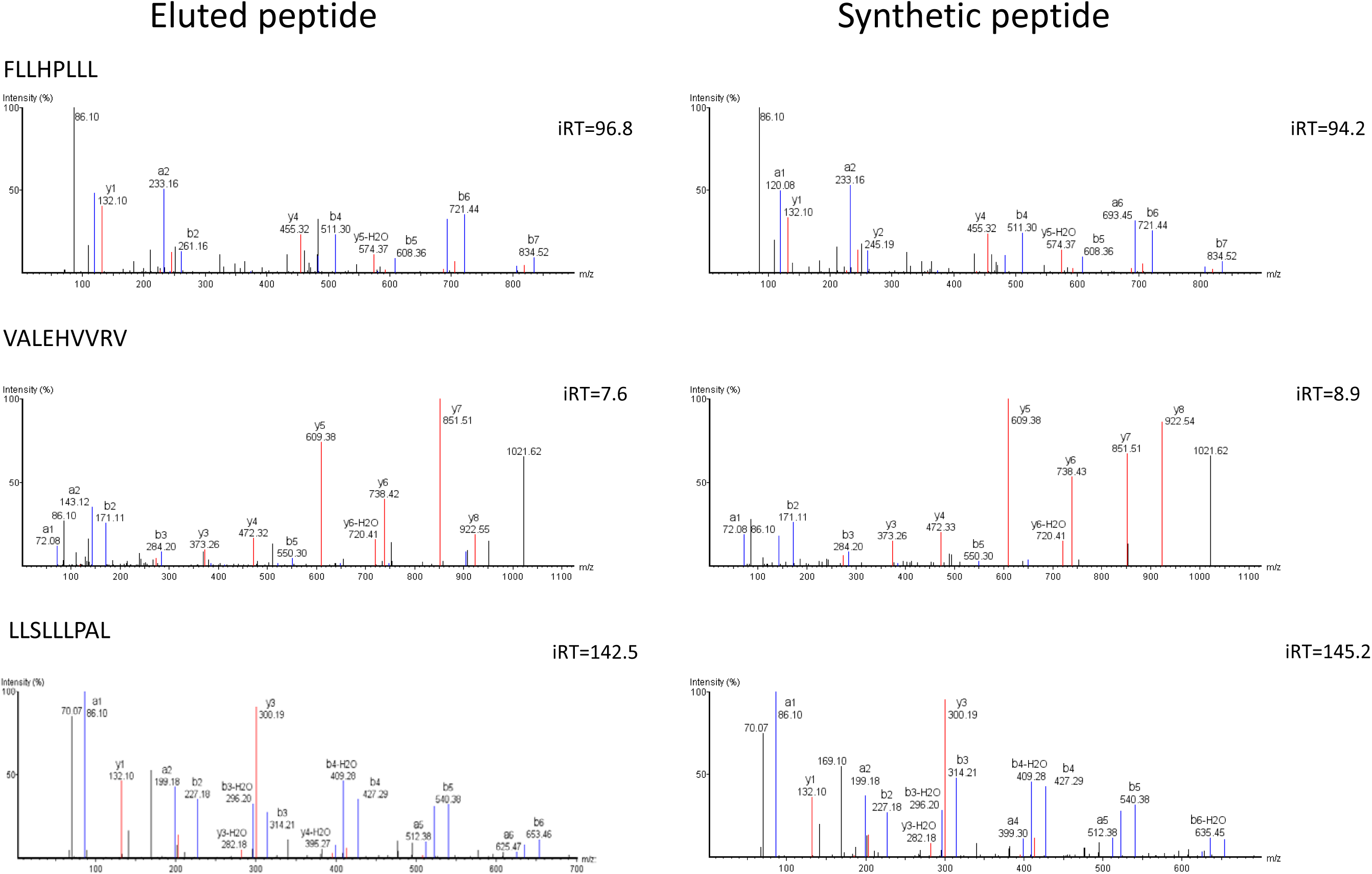
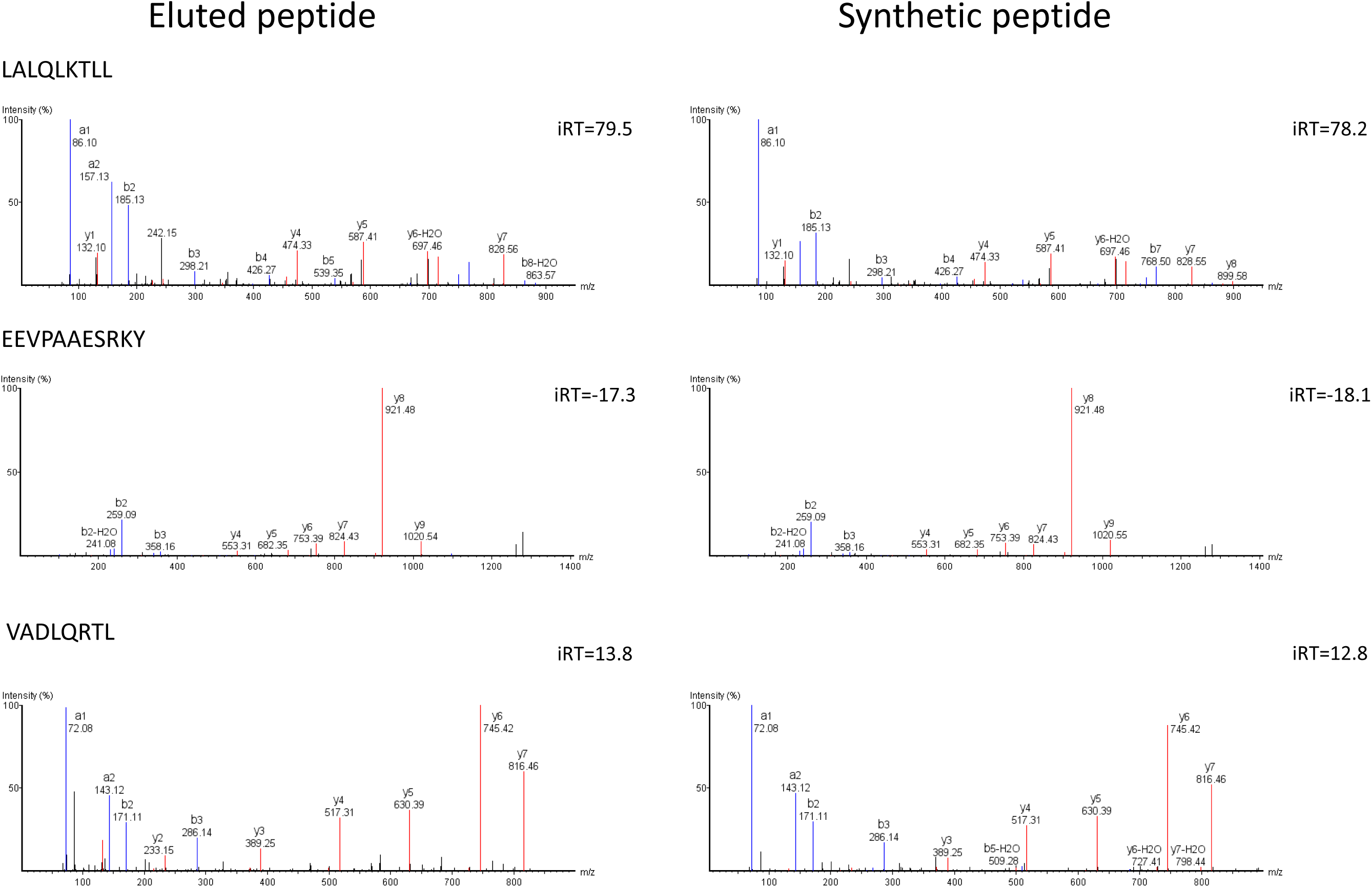
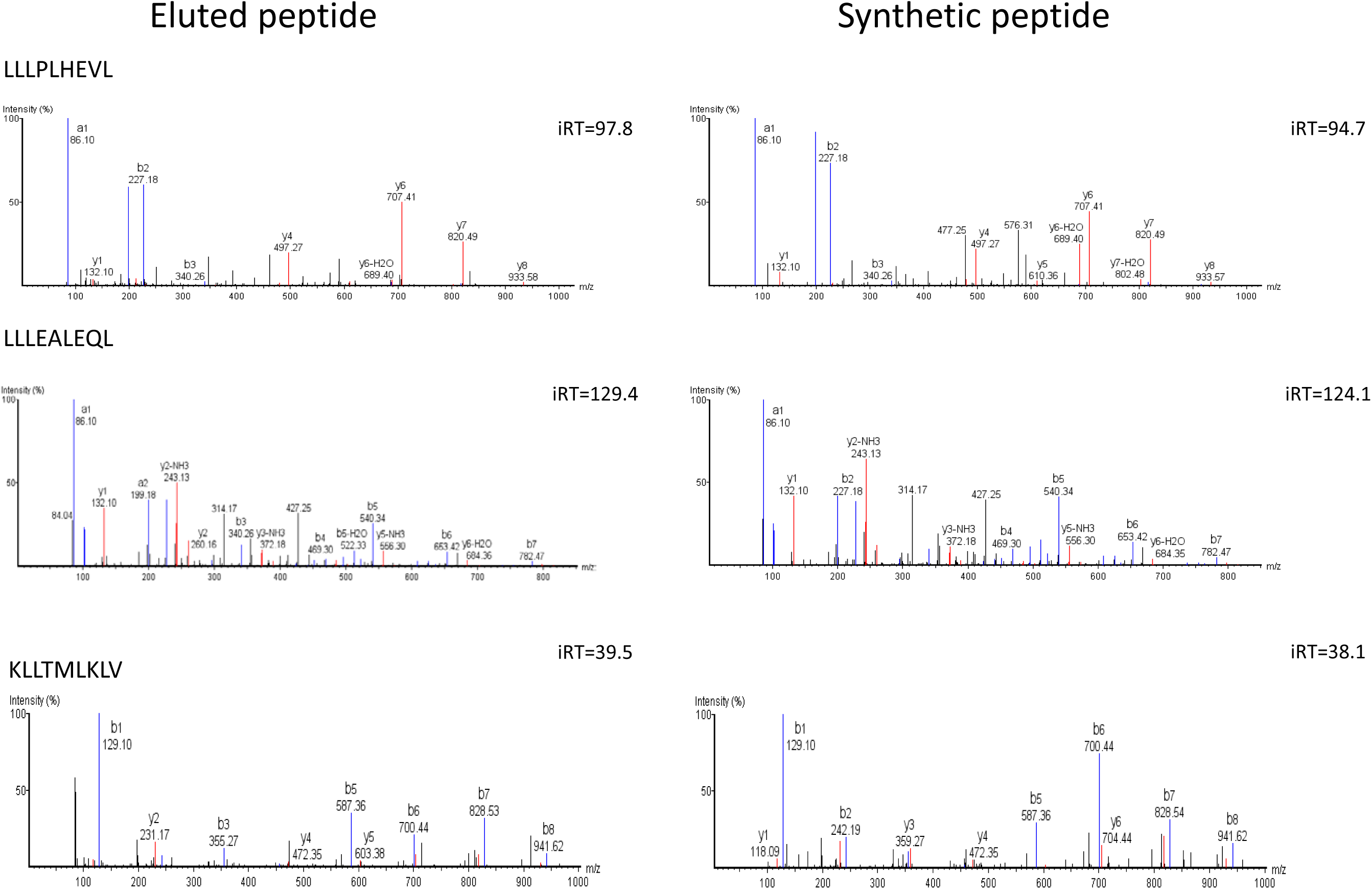
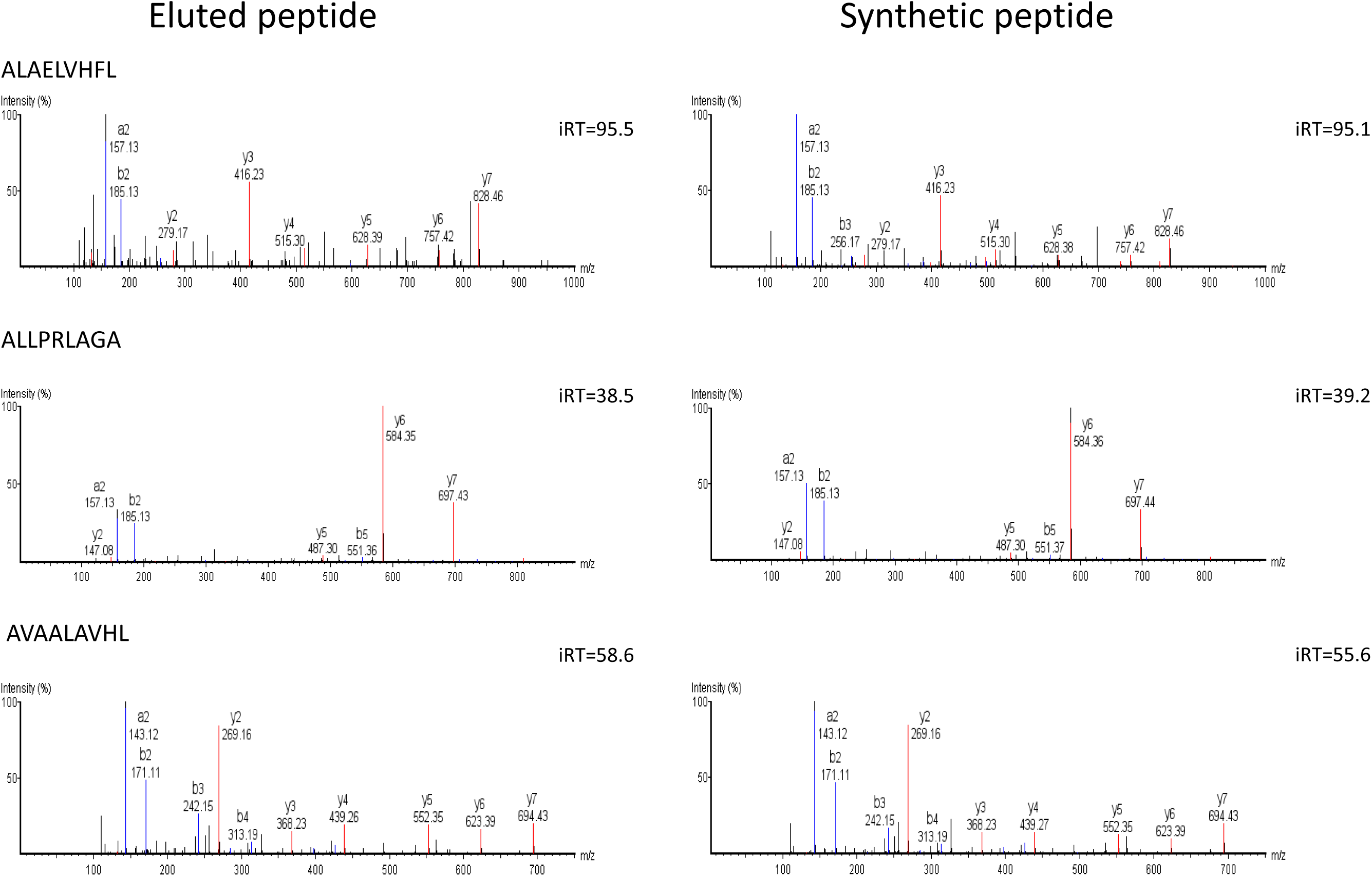
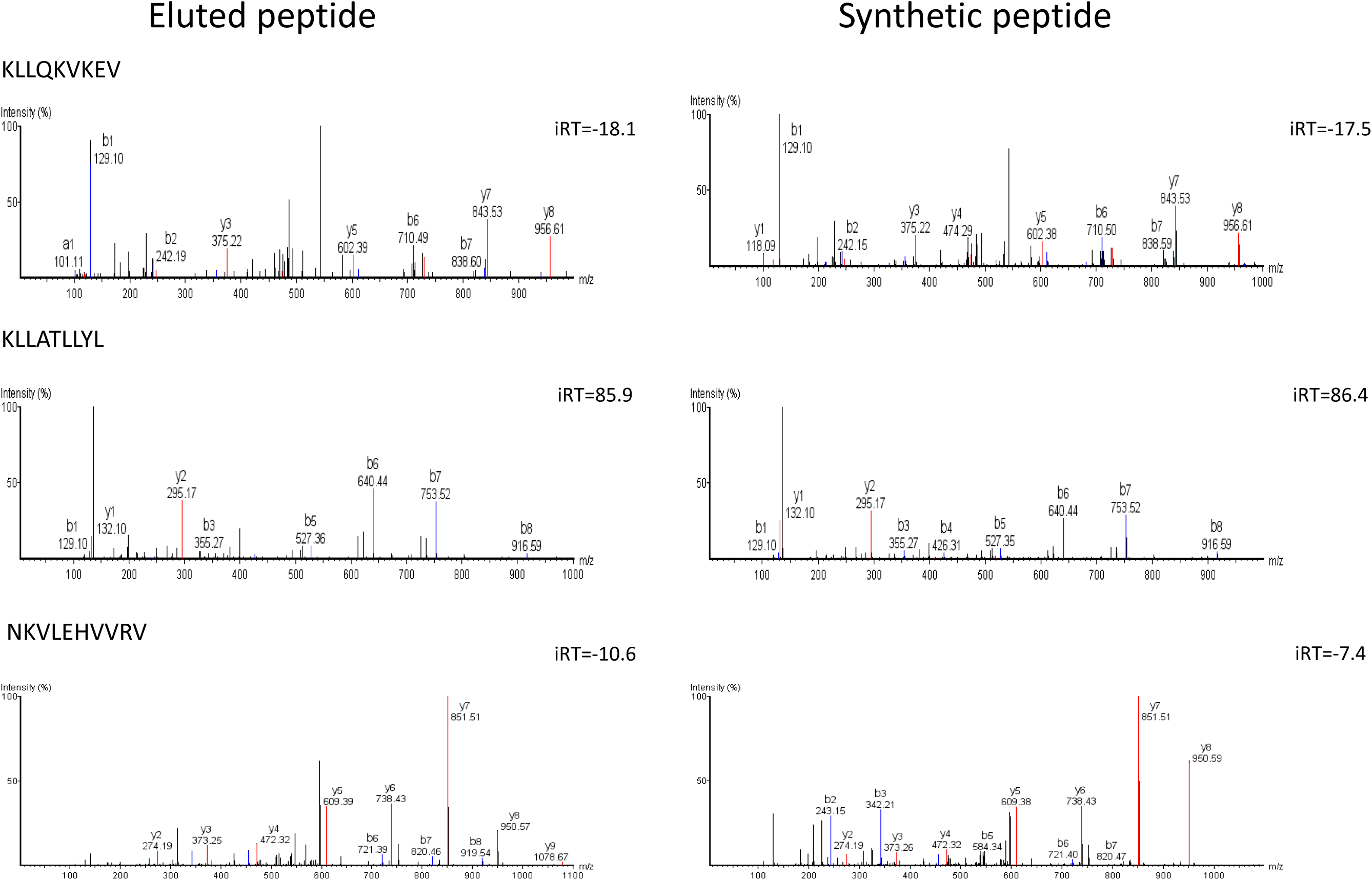
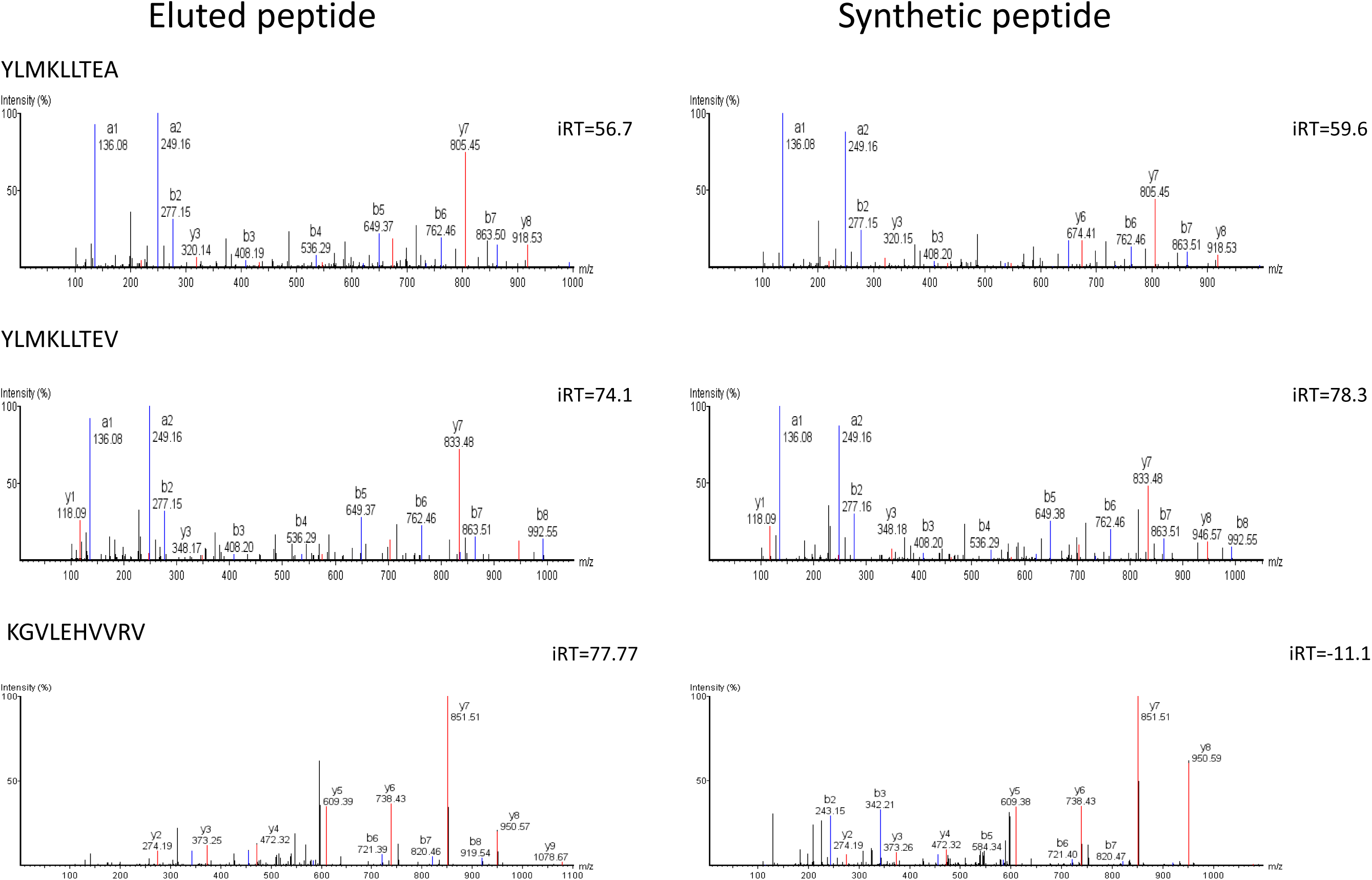
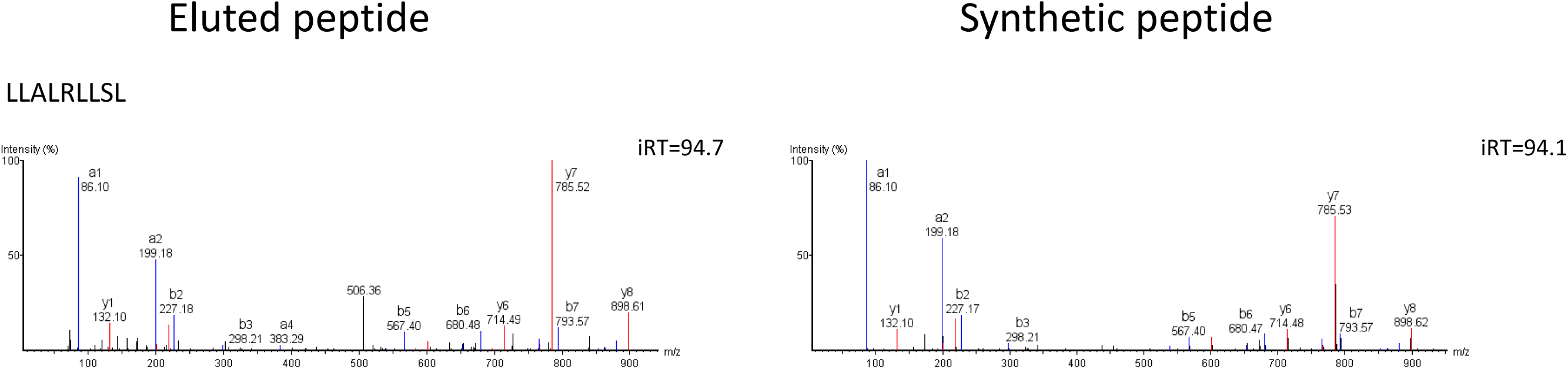
Comparison of MS2 spectra and iRT value of 28 *cis*-spliced peptides versus their corresponding synthetic version.

**Figure S9.**
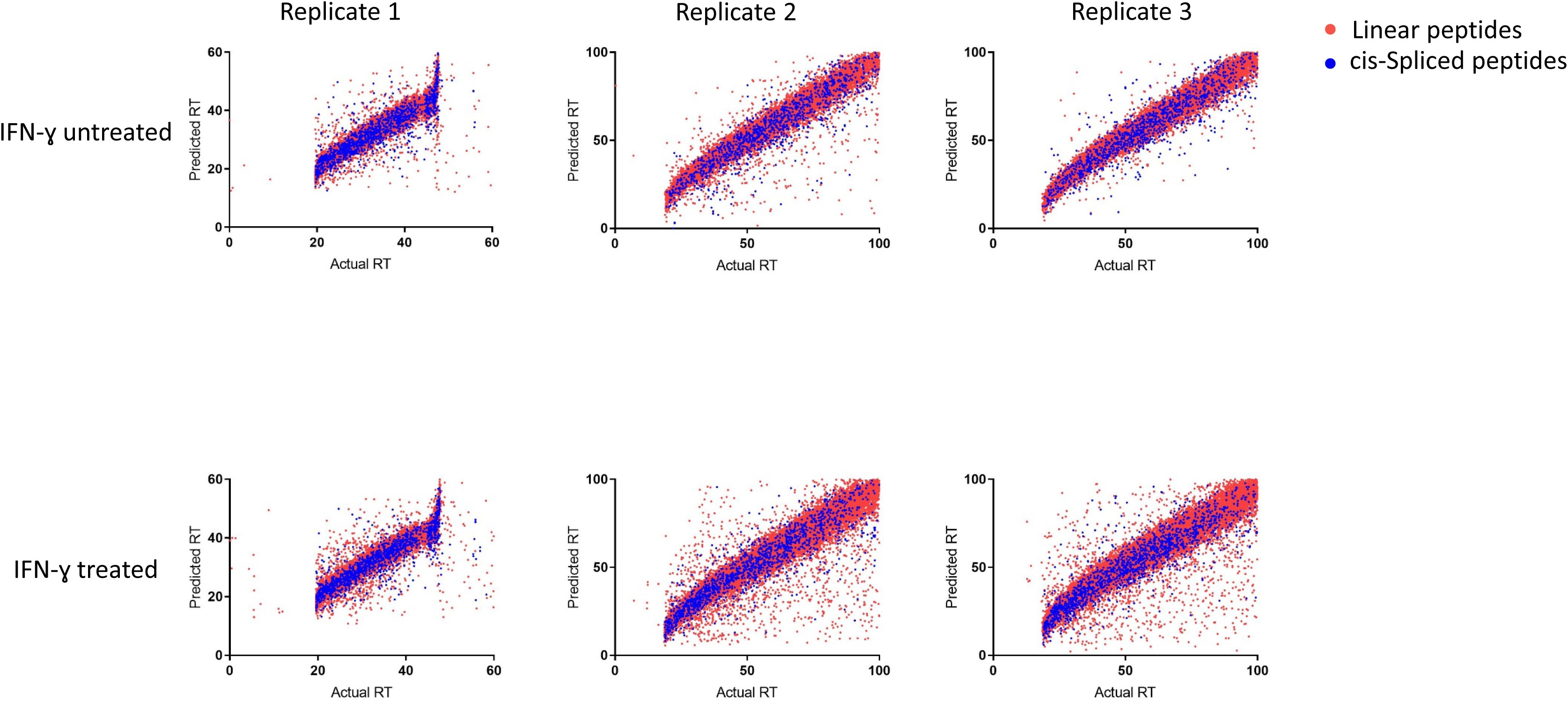
Comparison of predicted vs real retention time of linear and *cis*-spliced peptides by using GPTime tool (55). There was not a difference between linear and spliced peptide in this analysis in any of datasets.

## Supplementary Tables

Table S1. Identified Linear and *cis*-spliced peptides. Sheet 1; all 8-12 mer identified peptides and their presence in other cell lines or reported before (note that in spliced peptides, L stands for both leucine and isoleucine.). Sheet2; Shared peptides between six LM-MEL-44 samples. Sheet 3. List of LM-MEL-44 peptides presented in each condition. Sheet 4-13 raw export of PEAKS X software for each dataset.

Table S2. Binding prediction for identified linear and *cis*-spliced peptides from LM-MEL-44 cell line

Table S3. Predicted mutated peptides based on LM-MEL-44 sequencing data

Table S4. Melanoma associated antigens derived linear and *cis*-spliced peptides

Table S5. Selected linear peptides used for immunogenicity assay

Table S6: Presence of immunogenic linear and spliced sample across different cell lines

